# Embedding single-cell experimental conditions to reveal manifold structure of cancer drug perturbation effects

**DOI:** 10.1101/455436

**Authors:** William S. Chen, Nevena Zivanovic, David van Dijk, Guy Wolf, Bernd Bodenmiller, Smita Krishnaswamy

**Affiliations:** Department of Genetics, Yale School of Medicine; Yale University, New Haven, CT, USA; Applied Mathematics Program; Yale University, New Haven, CT, USA; Department of Computer Science; Yale University, New Haven, CT, USA; Institute of Molecular Life Sciences, University of Zurich, Zurich, Switzerland

## Abstract

Previously, the effect of a drug on a cell population was measured based on simple metrics such as cell viability. However, as single-cell technologies are becoming more advanced, drug screen experiments can now be conducted with more complex readouts such as gene expression profiles of individual cells. The increasing complexity of measurements from these multi-sample experiments calls for more sophisticated analytical approaches than are currently available. We developed a novel method called *PhEMD (Phenotypic Earth Mover’s Distance)* and show that it can be used to embed the space of drug perturbations on the basis of the drugs’ effects on cell populations. When testing PhEMD on a newly-generated, 300-sample CyTOF kinase inhibition screen experiment, we find that the state space of the perturbation conditions is surprisingly low-dimensional and that the network of drugs demonstrates manifold structure. We show that because of the fairly simple manifold geometry of the 300 samples, we can accurately capture the full range of drug effects using a dictionary of only 30 experimental conditions. We also show that new drugs can be added to our PhEMD embedding using similarities inferred from other characterizations of drugs using a technique called Nystrom extension. Our findings suggest that large-scale drug screens can be conducted by measuring only a small fraction of the drugs using the most expensive high-throughput single-cell technologies—the effects of other drugs may be inferred by mapping and extending the perturbation space. We additionally show that PhEMD can be useful for analyzing other types of single-cell samples, such as patient tumor biopsies, by mapping the patient state space in a similar way as the drug state space. We demonstrate that PhEMD is scalable, compatible with leading batch effect correction techniques, and generalizable to multiple experimental designs. Altogether, our analyses suggest that PhEMD may facilitate drug discovery efforts and help uncover the network geometry of a collection of single-cell samples.

## 1. Introduction

Single-cell data are now starting to be collected in large volumes and across numerous experimental conditions in order to characterize libraries of drugs, pools of CRISPR knockdowns, or groups of patients undergoing clinical trials. In such settings, the goal is to characterize the state space of the experimental variable, where the experimental variable may represent the specific perturbation condition or patient state. Put another way, the most interesting question in such experiments is often: *How do perturbation conditions (or patients) in a large set relate to one another, and in which ways do they differ?* In our study, we demonstrate that the state space of experimental variables may be embedded such that experimental conditions can be visualized, clustered and interpreted with respect to several intrinsic axes of variability. Our mapping technique is called *PhEMD*, or *Phenotypic EMD*.

PhEMD approaches the problem of studying the state space of an experimental variable by first solving the problem of comparing (i.e., computing a distance between) two clouds of points, wherein each point represents a single cell and each point-cloud represents a distinct multicellular experimental condition. Existing methods that aim to solve this problem include cellAlign [1] and sc-UniFrac [2]. However, both of these methods face significant limitations. cellAlign assumes all cells in an experimental condition lie on an unbranched trajectory, thus ignoring the intrinsic structure of the data and limiting the method’s utility to analyzing simple, unbranched transition processes. sc-UniFrac faces scalability issues and offers limited biological insights, as it does not reveal the *source* of differences between cell populations when performing pairwise comparisons of multiple experimental conditions (Supplementary Note 5). In light of these limitations, we developed a novel, scalable approach to comparing single-cell experimental conditions. PhEMD utilizes the Wasserstein metric [3], also known as Earth Mover’s Distance (EMD) [4] or optimal transport [5], to compare two experimental conditions. Intuitively, this distance measures the amount of energy an earth-moving vehicle would have to exert in order to transform one topographic landscape (e.g., distribution of dirt) into another landscape. This notion naturally lends itself to biological settings, where the “dirt” represents cells and the topographic landscape represents a distribution of cells across a range of cell subtypes (i.e., single-cell experimental condition). One can then interpret the “distance” between two experimental conditions as the “effort” required to transform the overall cell population of one condition to that of the other. In the present study, we use this model as a starting point and focus on computing a biologically-meaningful EMD that can be used to organize and characterize multi-sample single-cell data.

We demonstrate that once an intelligent distance between two single-cell experimental conditions is derived, an experimental variable state space embedding can be constructed by computing distances between each pair of experimental conditions. The distance matrix can be used to find new coordinates in which the experimental conditions can be embedded, visualized and clustered. We explore the properties of this final experimental variable state space embedding in depth and show that it is useful for relating all measured experimental conditions to one another simultaneously. We also show that such embeddings can be extended with additional data sources to include experimental conditions not directly measured with single-cell technologies, thus potentially reducing experimental burden when performing drug screens.

To demonstrate the utility of PhEMD, we used PhEMD to evaluate several systems including a new large perturbation screen we performed on breast cancer cells undergoing TGF-β-induced epithelial-to-mesenchymal transition (EMT), measured at single-cell resolution with mass cytometry. EMT is a process that is thought to play a role in cancer metastasis, whereby polarized epithelial cells within a local tumor undergo specific biochemical changes that result in cells with increased migratory capacity, invasiveness, and other characteristics consistent with the mesenchymal phenotype [6]. In our experiment, each perturbation condition consisted of cells from the Py2T breast cancer cell line stimulated simultaneously with TGF-β (to undergo EMT) and a unique kinase inhibitor, with the ultimate goal being to compare the effects of different inhibitors on our model EMT system. Using PhEMD, we embedded the space of the kinase inhibitors themselves and found that drug inhibition space was low-dimensional, with the drugs collectively acting along just three axes of variation as assessed through intrinsic dimensionality analysis. Further, we showed that due to the low-dimensional manifold geometry of the kinase inhibition space, just a select 10% of the inhibition conditions were needed to learn the geometry; the effects of the remaining kinase inhibitors could be imputed in relation to these.

In our drug-screen experiment, we found that some drugs effectively inhibited EMT and resulted primarily in epithelial cells despite chronic stimulation with TGF-β. Other drugs had little effect on EMT and resulted in the full spectrum of epithelial, transitional and mesenchymal cells. Still other drugs preferentially selected for apoptotic or other unique subpopulations of cells. We validated the drug-effect findings of our drug-screen experiment by showing that they were consistent with the drug-effect findings of a previously published study that profiled the drug-target binding specificities of several of the same drugs as ours. We specifically showed that the drug-drug relationships learned from our PhEMD analysis could be used to infer the results of the prior experiment (and vice versa). In doing so, we also demonstrated that the results of PhEMD could be combined with other datasets and data types to predict the phenotypes of samples not directly profiled by single-cell technologies.

To highlight generalizability of the PhEMD embedding approach, we performed analogous analyses on three additional datasets – one generated dataset with known ground-truth structure, one collection of 17 melanoma samples (scRNA-seq), and another of 75 clear-cell renal cell carcinoma samples (mass cytometry). In all experiments, the PhEMD embeddings revealed that the state space of the experimental variables could be modeled as a continuous manifold, often with biologically-interpretable dimensions. For example, in the melanoma and renal cell carcinoma datasets, the PhEMD manifolds highlighted the heterogeneity in patient samples with respect to tumor-infiltrating immune cells, showing the potential utility of PhEMD for disease subtyping. Collectively, our varied analyses demonstrate PhEMD’s wide applicability to numerous single-cell experiments.

## 2. Results

### 2.1 Overview of PhEMD

PhEMD is a method for embedding the state space of an experimental variable, in which each instance of the variable is measured with single-cell resolution. For example, when the experimental variable is the chemotherapy agent (drug) applied to cancer cells, each instance of the variable (i.e., specific drug condition) can be represented as the cancer cells stimulated with the drug, profiled using mass cytometry or single-cell RNA sequencing. Deriving an embedding of the drugs is useful for a variety of applications. These include visualizing the drug space to reveal the perturbation-effect landscape of the cancer, clustering drugs to identify treatments that induce similar responses in cancer cells, and modeling trajectories to reveal the main axes along which drugs vary with respect to their induced cellular response. Deriving an embedding involves finding a coordinate system (axes of variability) such that each point represents a drug and the distance between the points represents the dissimilarity between drugs with respect to induced cellular response. PhEMD derives such an embedding using the following general steps (Figure 1b):

**Figure 1:**
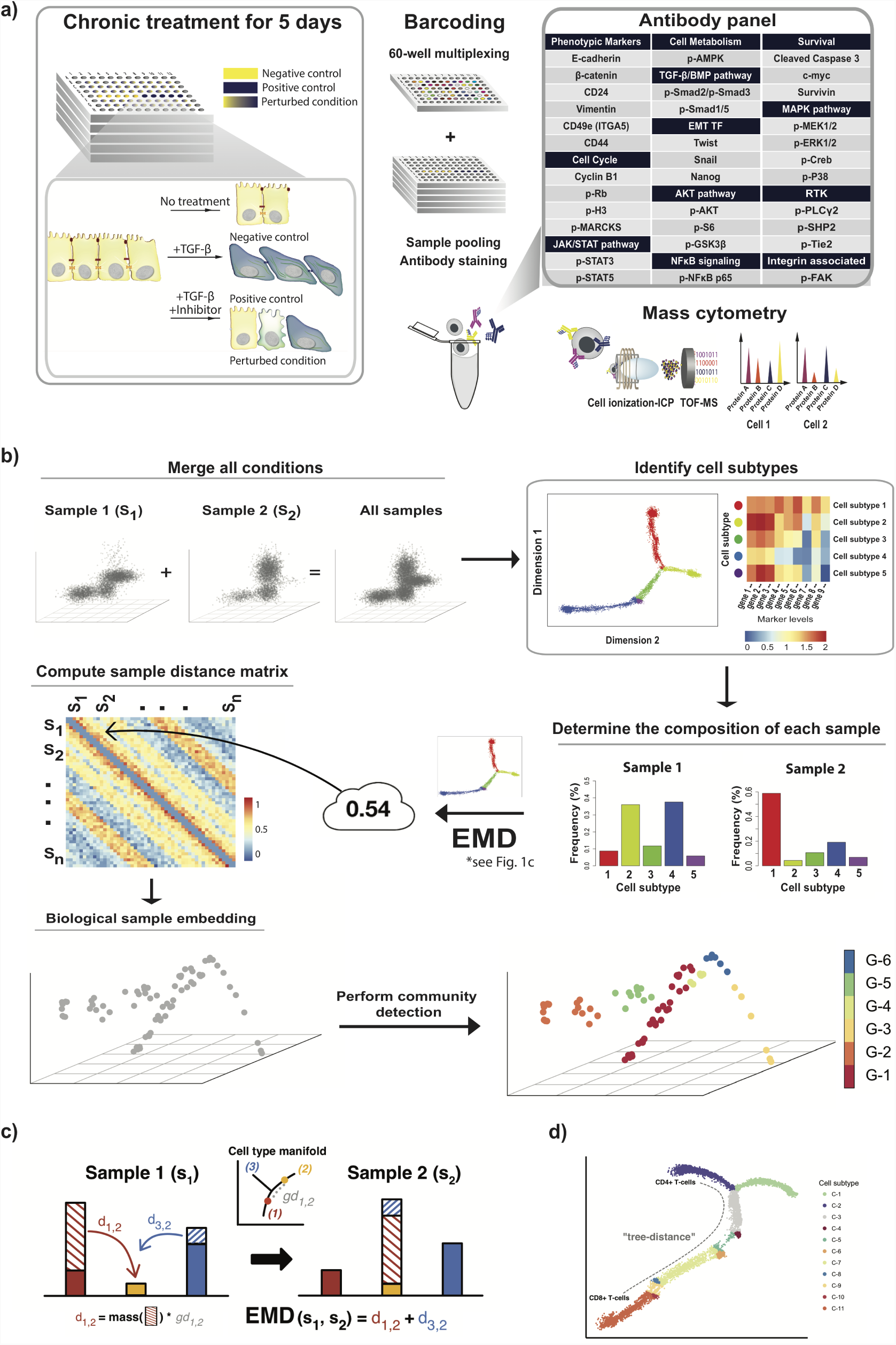
a) Experimental design for measuring perturbation effects of small molecule inhibitors on EMT. b) Flow diagram outlining the sequential steps performed in the PhEMD analysis pipeline. First, single cell measurements from all biological samples are aggregated. Next, unique cell subtypes and branched manifold structure are identified using Monocle 2. Then, deconvolution is performed to determine the composition of each unique biological sample (heterogeneous cell population) based on relative frequency of different cell subtypes. Earth Mover’s Distance is used as the measure of dissimilarity between two single-cell samples and is computed for each unique pair of samples to generate a distance matrix D. D is then used to generate an embedding using a diffusion map approach, and the resulting embedding can be visualized in low-dimensional space. Groups of similar samples are identified by performing community detection (e.g. hierarchical clustering) on D. c) Schematic of the EMD computation. The “completeness” of EMD as a distance metric is due to the fact that EMD takes into account both the differences in heights of matching bins and the intrinsic similarity of bins, as highlighted by the visual. d) Visual representation of “ground-distance” or “tree-distance” between cell subtypes, defined as the pseudotime distance between the centroids of the cell subtypes’ respective clusters. Tree-distance between subtypes C-2 and C-11 can be thought of intuitively as the length of the dotted path drawn in grey.

1. Compute a distance between each pair of single-cell samples (representing two settings of an experimental variable) by the following:
  a. Organize *cells* in the combined individual samples into a phenotypic tree.
  b. Cluster cells on this phenotypic tree and then represent each of the individual samples as proportions of these clusters.
  c. Compute the distance between two samples using Earth Mover’s Distance. This is the optimal transport distance for moving cells from the way they are proportioned between clusters in one sample to the way they are proportioned in the other. The transport distance increases with the number of cells moved and the dissimilarity of cell subtypes between samples, as it penalizes not only how many cells need to be moved but also how “far” each cell is moved along the phenotypic tree. This penalty involving dissimilarity of cell subtypes is known as the *ground distance* (Supplementary Note 4).
2. Take the derived distance matrix from the previous step, convert the distances into affinities using a Gaussian kernel, and Markov-normalize (row-normalize) this kernel to obtain probabilities rather than affinities in each row.
3. Eigendecompose the matrix from the previous step to find the coordinates of the embedding.

Pseudocode for the PhEMD algorithm is shown below and additional details can be found in the Online Methods section.

#### Algorithm 1

**Pseudocode for the PhEMD analytical approach**

1: **procedure** PHEMD(*single.cell.data*)

2: ⊲Define cell subtypes

3:*data.al*← aggregateCells_all_samples(*single.cell.data*)

4:*cellT ype.embedding, cellT ype.assignments* ← Monocle2(*data.all*)^*\**^

5:visualize_cell_embedding(*cellType.embedding*)

6:visualize_heatmap(*cellType.assignments, data.all*)

7:

8: ⊲Compare samples based on cell subtype relative frequencies

9:*cellT ype.freq*←deconvolute samples(*data.all, cellT ype.assignments*)

10:**for** each pair of samples *si, sj* **do**

11:*Dists*[*i, j*] ← EMD(*cellType.signatures, cellT ype.freq*[*i*], *cellT ype.freq*[*j*])

12:*biological.sample.embedding*←DiffusionMap(*Dists*)

13:*biological.sample.groups*←HierarchicalClustering(*Dists*)

14:visualize_sample_embedding(*biological.sample.embedding*)

15:visualize_sample_cellTypeFrequencies(*cellT ype.freq*)

*See Supplementary Note 7

### 2.2 PhEMD correctly reconstructs cell-state geometry and biological sample embeddings on high-dimensional data with known ground-truth structure

We first applied PhEMD to generated data with a known ground-truth tree structure to determine whether it could accurately model both the cell-state and biological-sample embeddings. The simulated cells lay on a continuous branched trajectory, where branches represented concurrent increases or decreases in multiple distinct dimensions in 60-dimensional space [7]. To simulate multiple biological samples, we varied the distribution of point density across branches of the cell-state tree between samples (Online Methods).

PhEMD was able to construct the correct cell-level tree structure using Monocle 2 (Supplementary Figure 3a). The inter-sample EMD-based comparison and resulting embedding were also found to be accurate (Supplementary Figure 3b). This accuracy was assessed two-fold. First, we examined those samples in which a large number of points were concentrated in a single branch. We found that samples with point density concentrated in branches close to one another on the cellstate tree (e.g. Samples X and Y) tended to map to regions close to one another on the biological-sample manifold compared to samples with point density concentrated in branches far from one another on the cell-state tree (e.g. Samples X and Z). Next, we examined Samples A–K: samples in which point density was modulated so that Sample A had points mostly in the arbitrary “starting” state of the tree, Sample K had points mostly in an arbitrary “terminal” state, and samples B through J had progressively fewer points in the “starting” state and more points in the “terminal” state. We found that in the final biological sample embedding, Samples A–K appropriately formed a trajectory and were ordered based on their intra-sample relative proportions of “starting state” to “terminal state” points.

**Figure 3:**
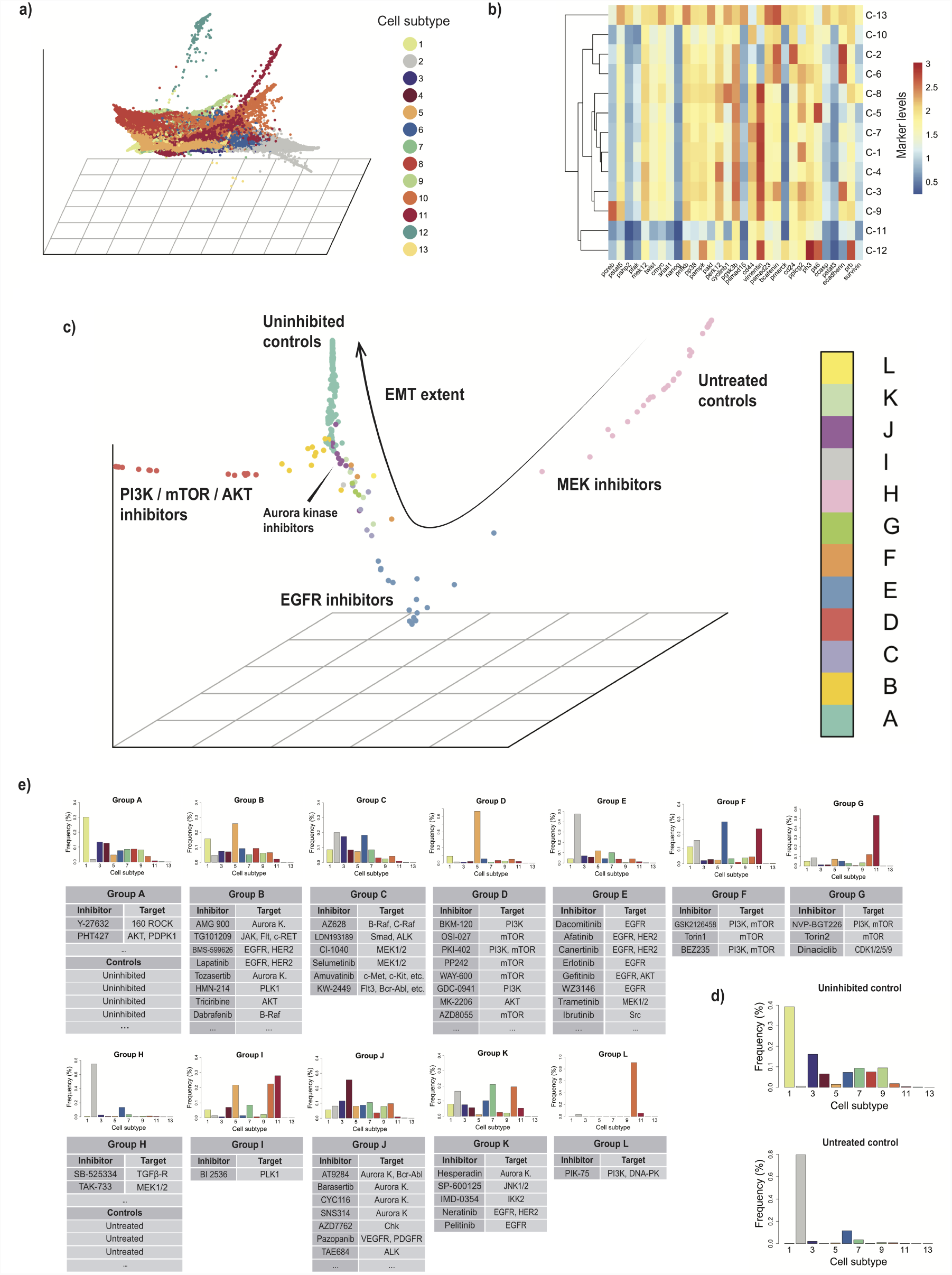
a) PHATE embedding of cells from all 300 experimental conditions, colored by Louvain-based cell subtype. b) Heatmap representing log_2_ protein expression levels for each cell subpopulation representing its respective cell subtype. c) Embedding of control and drug-inhibited conditions, colored by inhibitor clusters assigned by hierarchical clustering. d) Distribution of cells across cell subtypes for uninhibited (TGFβ-only) and untreated controls. e) Individual inhibitors assigned to each inhibitor group. Histograms represent bin-wise mean of relative frequency of each cell subtype for all inhibitors in a given group.

### 2.3 Effect of drug perturbations on the EMT landscape in breast cancer

To study key regulators of epithelial-to-mesenchymal transition (EMT) in breast cancer, we performed a drug screen consisting of 300 inhibition and control conditions, collectively inhibiting over 100 unique protein targets (mostly kinases) in murine breast cancer cells undergoing TGFβ-induced EMT (Figure 1a; Online Methods; Supplementary Table 1). These samples collectively contained over 1.7 million cells measured in a total of five mass cytometry runs. Time-of-flight mass cytometry (CyTOF) was used on day 5 of cell culture to measure the concurrent expression of 33 protein markers in each cell (Supplementary Table 2). PhEMD was subsequently used to model both the cell-state transition process and the drug perturbation manifold (Figure 2). To avoid the potential confounding effects of batch effect (especially when characterizing the EMT cell state space), we first analyzed inhibition and control conditions from a single experimental run. We subsequently performed batch correction and analyzed all samples across all plates simultaneously. Three biological replicates were analyzed to demonstrate reproducibility of results (Supplementary Figure 4).

**Figure 2:**
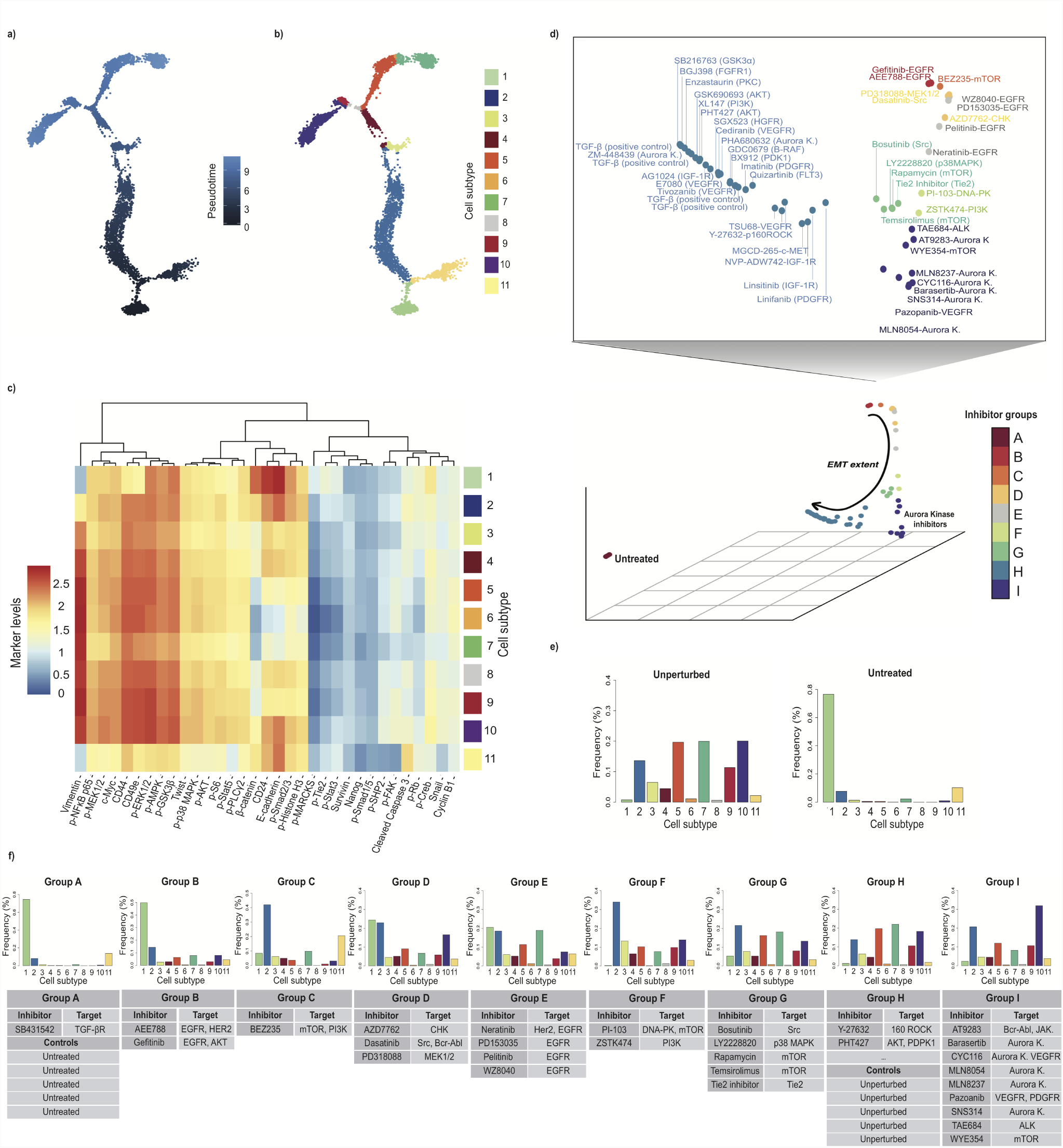
Monocle 2 embedding of cells from all conditions of a single CyTOF run representing perturbed EMT cell state landscape, colored by a) pseudotime and b) cell subtype. Since all cells were known to have been derived from an epithelial population, the epithelial cell state was defined as the “starting state” of the tree-based model when performing pseudotime ordering of cells. c) Heatmap of log_2_ protein expression levels for each subpopulation of cells representing a distinct cell subtype. d) Embedding of drug inhibitors, colored by clusters assigned by hierarchical clustering. e) Distribution of cells across cell subtypes for unperturbed (TGFβ-only) and untreated controls f) Individual inhibitors assigned to each inhibitor group. Histograms represent bin-wise mean of relative frequency of each cell subtype for all inhibitors in a given group.

**Figure 4:**
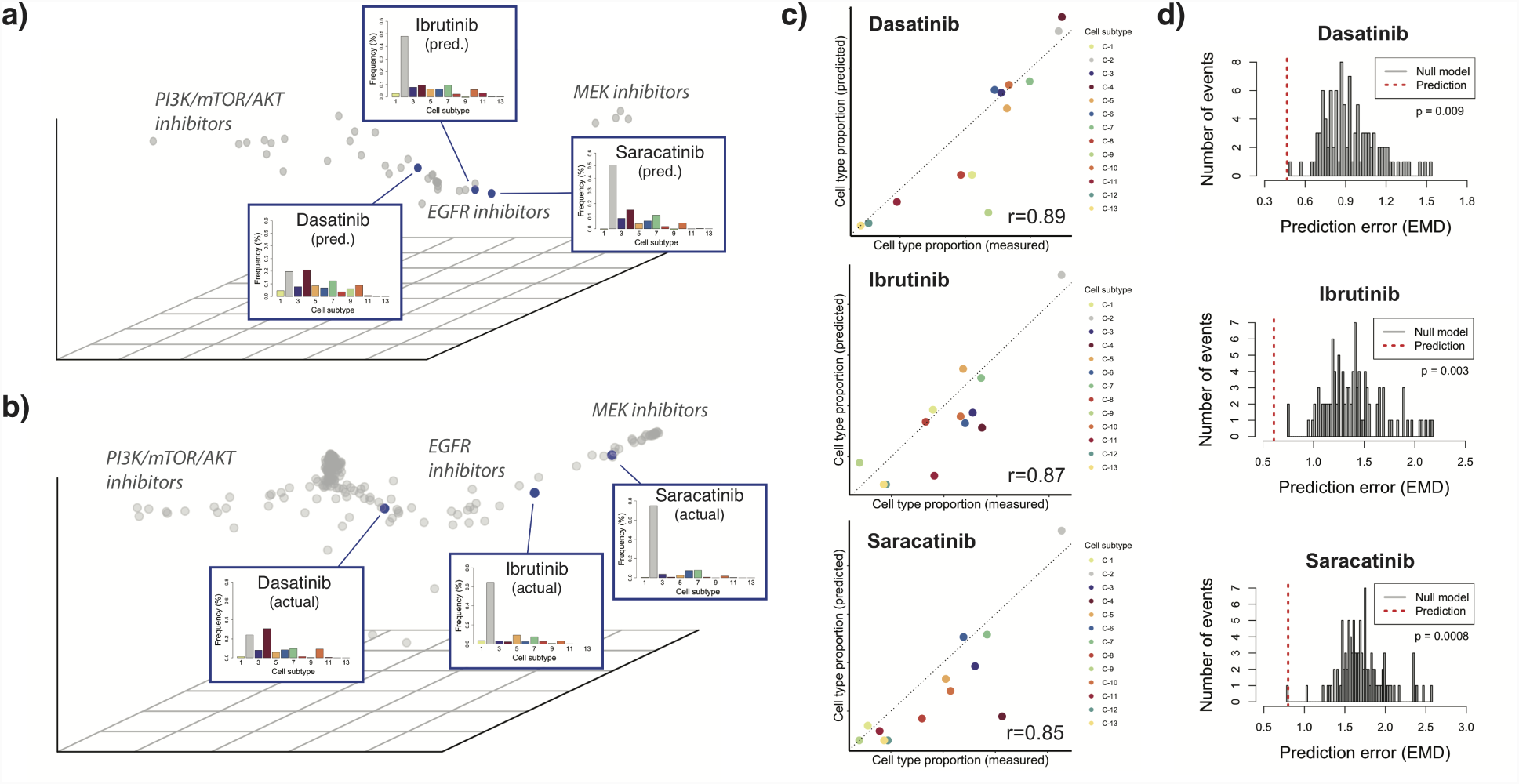
a) Nystrom extension embedding showing predicted effect of 3 selected inhibitors (dasatinib, ibrutinib, saracatinib) on EMT relatively to other measured inhibitors. b) PhEMD diffusion map embedding showing measured effects of 3 selected inhibitors on EMT. c) Scatterplot showing correlation between predicted vs. measured cell subtype relative frequencies for 3 selected inhibitors. Each point represents a cell subtype, colored using the same coloring scheme as histogram bars in Figures 6a-b. *x* = *y* line is shown in dotted grey. d) Histogram showing distribution of prediction error for null model. Dotted red line represents prediction error for actual prediction (i.e. alternative model).

#### 2.3.1 Analyzing EMT perturbations measured in a single CyTOF experimental run

##### 2.3.1.1 Cell subtype definition via tree-based clustering

To most accurately model the cell state space, we first characterized a subset of 60 inhibition and control conditions that were measured in the same CyTOF run. By design, we knew that all cells undergoing EMT lay on a continuous manifold, as they were all derived from the same relatively homogeneous epithelial population (i.e., same cell line). Thus, a continuous branched trajectory (i.e. tree) as modeled by Monocle 2 was ideal to relate the cell subtypes in our EMT experiment. Our tree-based model of the cell-state space identified 11 unique cell subtypes across all unperturbed and perturbed EMT samples (Figure 2). These included the starting epithelial subtype (C-1), main mesenchymal subtype (C-7), and transitional subtypes on the major EMT-axis (C-2 through C-6). C-1 was characterized by the following expression pattern: E-cadherin^(hi)^ β-catenin^(hi)^ CD24^(hi)^ p-CREB^(hi)^ vimentin^(lo)^ CD44^(lo)^. C-6 and C-7 had roughly the opposite expression profile with respect to the markers described above (Figure 2c). E-cadherin is the hallmark cell adhesion marker of epithelial cells [8], and vimentin and CD44 are known mesenchymal markers involved in cell migration [8–11]. Moreover, recent studies found high CD44:CD24 expression to be indicative of breast cancer cell invasiveness and an as an EMT endpoint, suggestive of mesenchymal properties [12–14]. Altogether, the subtypes identified by Monocle 2 are consistent with known epithelial and mesenchymal cell phenotypes, and the trajectory defined by subtypes C-1 through C-7 in our model represent the epithelial-to-mesenchymal transition process that one would expect to recover in our dataset.

In addition to modeling the main EMT trajectory, the Monocle 2 cell-state embedding identified two branches (sub-trajectories) off the main EMT axis. The proximal branch represented an epithelial subpopulation undergoing apoptosis (C-11), with high E-cadherin and cleaved caspase-3 expression. The second, more distal “hybrid EMT” branch was defined by cells with intermediate levels of E-cadherin and vimentin and increased expression of p-MEK1/2, p-ERK1/2, p-p38-MAPK, p-GSK-3β, and p-NFkB-p65 (C-8 through C-10). This unique cell population and our non-linear model of EMT were consistent with the recent discovery of “hybrid” cancer cells that co-express epithelial and mesenchymal markers (E+/M+) and simultaneously demonstrate both epithelial and mesenchymal properties [15–17]. By analyzing our single-cell data with Monocle 2, which applied no prior assumptions on the total number of branches in the cell-state embedding, we were able to uncover a more complex, continuous model of EMT than has been previously reported that captures the E+/M+ cell population of increasing clinical and biological interest.

##### 2.3.1.2 Constructing the EMD-based drug-inhibitor manifold

After modeling the EMT cell-state space with Monocle 2, we used PhEMD to derive a distance between drug inhibitors based on the relative distribution of cell density across the different cell sub-types identified above. Specifically, EMD was computed pairwise between inhibition conditions to construct a distance matrix (Online Methods). In this experiment, EMD represented the minimum “effort” required to transform one inhibition condition to another (conceptually equivalent to the total “effort” needed to move cells from relatively “overweight” parts of the branched, continuous, EMT cell-state tree to relatively “underweight” parts). After deriving the EMD between every pair of inhibition conditions, we effectively had a network of drug inhibition conditions, represented as an EMD-based distance matrix. This matrix could be embedded using dimensionality reduction techniques and subsequently visualized. We chose to use a diffusion map embedding of this distance matrix to capture continuous and non-linear relationships between samples.

To assess the innate complexity of our perturbation state space, we performed intrinsic dimensionality analysis on our network of 60 inhibition and control conditions. Using the maximum likelihood estimation approach, we found that the intrinsic dimensionality of our data was approximately two across a range of reasonable parameters (Online Methods; Supplementary Figure 5a). This suggested that our 60-sample PhEMD embedding could be reasonably represented and visualized in two dimensions.

**Figure 5:**
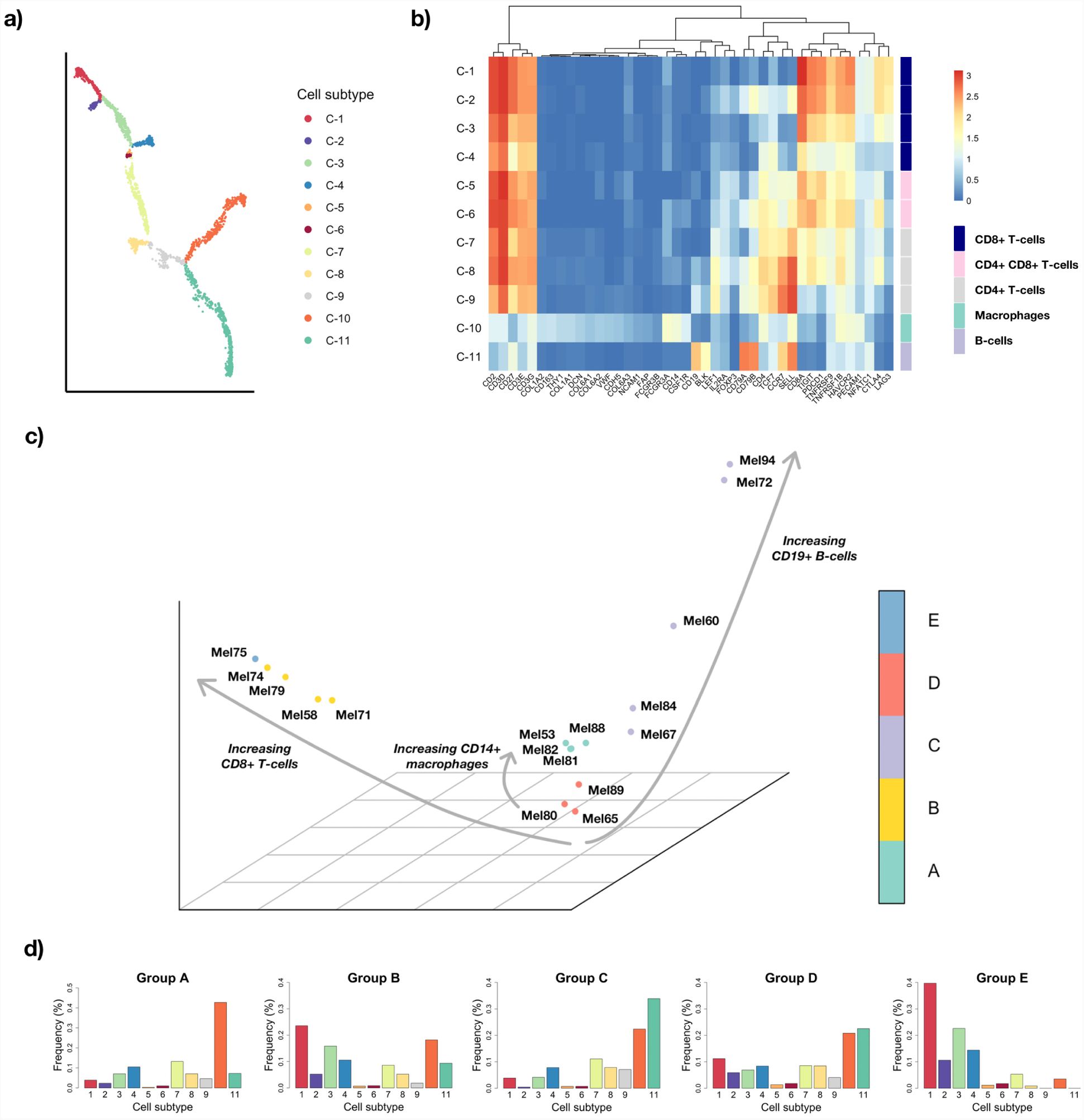
PhEMD applied to scRNA-sequencing data of 17 melanoma samples (non-tumor cells only) highlights heterogeneous immune response amongst different patients. a) Monocle 2 cell state embedding colored by cell subtype. b) Heatmap showing mean RNA expression values of each cluster, colored by a log_2_ scale. c) Diffusion map embedding of samples (colored by group assignment) revealing multiple trajectories that represent increasing relative frequency of selected cell populations. d) Summary histograms, each representing the bin-wise mean relative frequency of cell subtypes for all samples assigned to a given group.

##### 2.3.1.3 Clustering the drug-inhibitor manifold

The EMD-based embedding of drug inhibitors (constructed as described above) was then partitioned using hierarchical clustering. Note that once a geometry of experimental conditions (i.e., single-cell samples) was constructed as we did, the *samples* could be compared using tools similar to those typically used to compare individual *cells* (e.g. for cell clustering and trajectory modeling). Hierarchical clustering revealed clusters of inhibitors with similar net effects on EMT; inhibitors assigned to the same cluster were assumed to have similar effects on EMT. Moreover, by including “uninhibited” controls (samples in which TGF-β was applied to induce EMT in absence of any inhibitor) and “untreated” controls (samples in which neither TGF-β nor inhibitor was applied and no EMT was induced) in our experiment, we were able to identify inhibitors with notable effects on EMT. Those inhibition conditions that clustered with uninhibited controls likely had little to no effect on EMT, whereas those that clustered with untreated controls halted EMT strongly and likely at an early stage.

Our embedding of drug inhibitors revealed a manifold structure that highlighted the variable extent of EMT that had occurred in the different inhibition conditions (Figure 2d). Partitioning the embedding into nine clusters (Clusters A-I; Supplementary Figure 6, Supplementary Table 3), we found that Cluster A included the untreated controls and the TGF-β-receptor inhibitor condition, each of which consisted almost entirely of epithelial cells. These were the experimental conditions in which EMT was actually or effectively not induced. On the other hand, Cluster H included all five uninhibited control conditions and inhibitors ineffective at modulating EMT; inhibitors in this cluster were found to have mostly late transitional and mesenchymal cells. Clusters B through G included inhibitors that had generally decreasing strength with respect to halting EMT (Figure 2, Supplementary Note 9). The two EGFR inhibitors in Cluster B strongly inhibited EMT, as indicated by a marked predominance of epithelial cells at time of CyTOF measurement, and the five inhibitors in Cluster G each had a mixture of epithelial, transitional, and mesenchymal cells, with a predominance of mesenchymal cells. In general, small molecule inhibitors that had the same molecular target tended to cluster together, which was consistent with the intuitive notion that drugs with similar mechanisms of action would likely have similar net effects on a given cell population (e.g. Cluster B, Cluster I). However, we also noted that some inhibitors with the same reported primary target generated different resulting cell profiles and were clustered into different inhibitor clusters (e.g. Cluster B and Cluster E). This phenomenon may be due to differences in inhibitor potency and differences in off-target effects.

**Figure 6:**
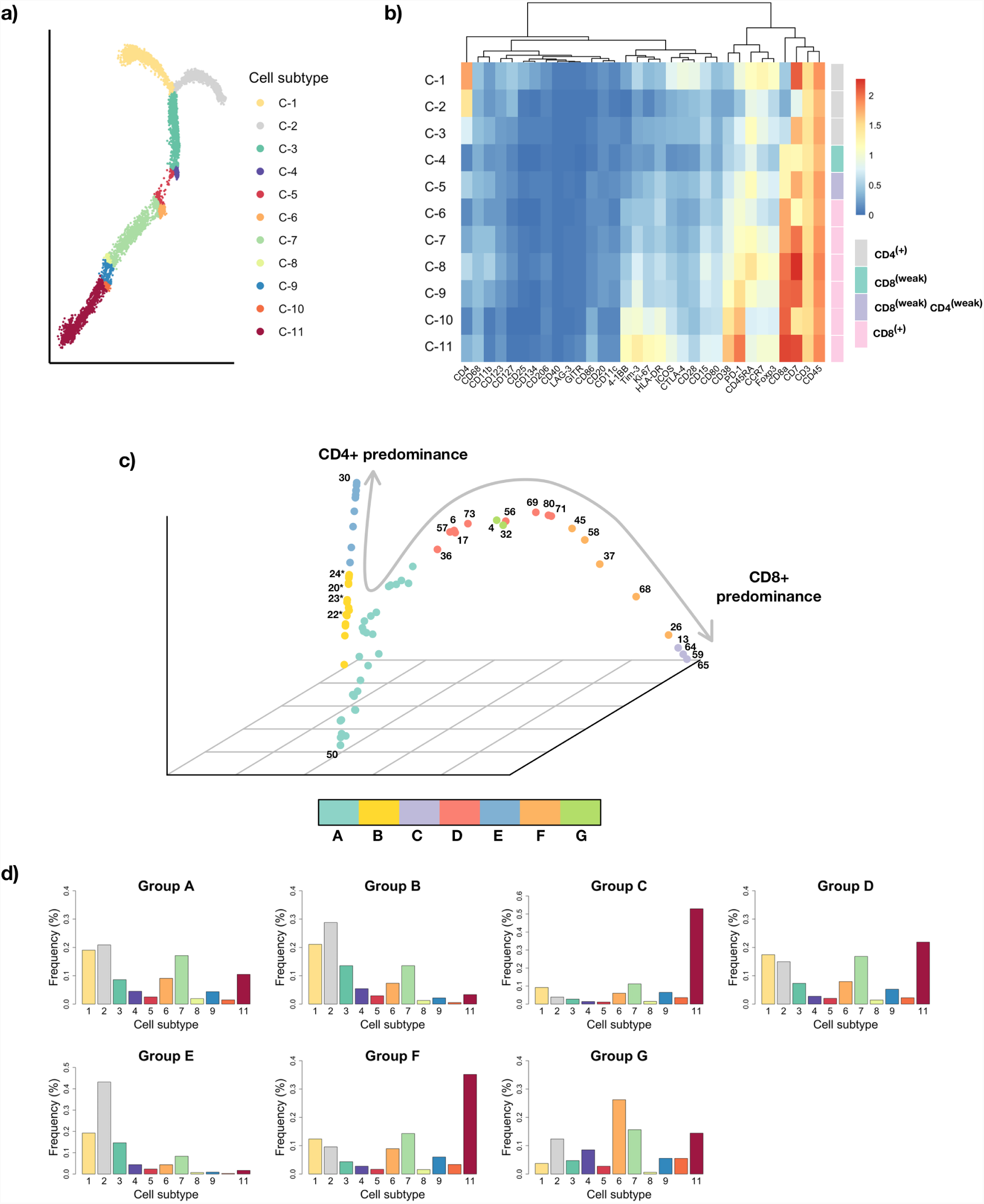
PhEMD applied to mass cytometry data of 75 ccRCC samples gated for T-cells. a) Monocle 2 embedding of T-cell manifold colored by 11 cell subtypes identified by the algorithm.b) Heatmap showing mean protein expression values of each cell subtype cluster, colored by a log_2_ scale of intensity. c) Diffusion map embedding of all tumors colored by tumor subgroup, which is defined by hierarchical clustering. Numbered labels represent sample IDs assigned in the original dataset, and numbers accompanied by asterisks denote healthy controls. d) Summary histograms, each representing the bin-wise mean relative frequency of cell subtypes for all samples assigned to a given group.

We found that inhibitors in Cluster I formed a small branch off the main EMT-extent trajectory in the inhibitor embedding (Figure 2d). These inhibitors consisted mostly of Aurora kinase inhibitors, and each demonstrated a cell profile characterized by a relatively high proportion of cells in the hybrid EMT trajectory (C-8 through C-10 in Figure 2a). Examining these results alongside measurements of cell yield in each inhibition condition (Supplementary Table 4), we attributed the relatively greater proportion of C-10 cells in the Aurora kinase inhibitors to preferential drug-induced death of other cell types. C-10 cells were not uniquely generated by Aurora kinase inhibition, as they were observed in other samples including the uninhibited EMT control conditions (Figure 2c) but appeared to have increased cell viability relative to other EMT cell types, especially in the setting of Aurora kinase inhibition (Supplementary Table 4).

#### 2.3.2 Single-cell inhibitor screen involving 300 inhibition and control conditions measured in five experimental runs

After analyzing results from a single experimental run in detail, we aimed to characterize our entire network of 300 inhibition and control conditions measured in a total of five mass cytometry runs. To correct for batch effect, we performed canonical correlation analysis (CCA) using the implementation included in the Seurat single-cell analysis suite [18]. Since Monocle 2 cannot be used with the batch-corrected output of CCA, we instead used the Louvain community detection algorithm included in Seurat to define cell subtypes. We related subtypes to one another based on their Euclidean distance in the aligned, dimensionality-reduced CCA space (Online Methods).

CCA was able to successfully correct for batch effect in our multi-run analysis (Supplementary Figure 7, Supplementary Note 10), and Louvain community detection identified 13 cell subtypes. Similarly to our single-experiment analysis of the EMT cell state space (Figure 2), Louvain community detection identified a mixture of epithelial (C-2, C-6), transitional (C-2, C-3, C-7 through C-11), mesenchymal (C-1, C-4, C-5, C-12), and apoptotic (C-13) cells (Figure 3). After defining cell subtypes, PhEMD effectively modeled a low-dimensional embedding of all 300 inhibition and control conditions, with an intrinsic dimensionality of three (Online Methods; Supplementary Figure 5b). PhEMD identified 12 clusters of inhibitors with similar effects on EMT (Supplementary Table 5). As could be expected, the PhEMD embedding and inhibitor clusters were similar to those of our single-plate analysis (Figure 2) while including a more comprehensive set of inhibitors with a collectively larger set of protein targets.

Cluster A was the largest and included all uninhibited controls. The remaining samples in this cluster were inhibition conditions that were deemed ineffective at influencing EMT, since they generated a cell subtype distribution closely resembling that of the uninhibited controls. In our final PhEMD embedding, we found that samples in Cluster A were tightly clustered in a particular region of the low-dimensional space – visually demonstrating the relatively high intra-group similarity (Figure 3). Cluster H was found at the other end of the PhEMD embedding and included all of the untreated control conditions (comprised almost exclusively of epithelial cells). Several MEK inhibitors, a Src inhibitor, and a TGF-β-receptor inhibitor were also included in this group, suggesting that they strongly halted EMT. Just as in our single-plate analysis, we found that many of the inhibitors in the PhEMD embedding formed a trajectory that roughly corresponded to the strength of EMT inhibition. In roughly decreasing strength, these included EGFR inhibitors (Cluster E), certain PI3K/mTOR inhibitors (Cluster F), and additional inhibitors in Clusters C, G, I, K, and L.

Just as in our single-batch analysis, we found that many Aurora kinase inhibitors formed a cluster (Cluster J) that was enriched in cell subtypes C-4, C-9, and C-10. These subtypes demonstrated a similar expression profile to the hybrid-EMT profile identified during single-plate analysis (C-8 through C-10 in Figure 3b). However, the most striking finding in this expanded drug screen was a prominent trajectory formed by Clusters B and D. Clusters B and D were enriched in cell subtype C-5, with Cluster D inhibitors inducing cell populations that were almost entirely comprised of C-5 cells. Of note, all of the Cluster D inhibitors targeted either PI3K, mTOR, or AKT – three related molecules in a well-characterized pathway.

Compared to the predominant mesenchymal subtype observed in the uninhibited controls (C-1), C-5 was comprised of slightly lower expression of most markers and markedly higher expression of phospho-S6 (Figure 3). This profile was consistent with a late-transitional or alternative-mesenchymal EMT subtype. Examining the cell yield of these inhibitors compared to the respective uninhibited control conditions in their respective batches, we found that the cell yield of the Cluster D inhibitors was on average 60% lower than the TGF-β-only controls (Supplementary Table 4). Based on these findings and a prior report suggesting that decreased phosphorylation of ribosomal protein S6 may be associated with sensitivity to certain targeted therapies [19], it is possible that the C-5 subtype may be relatively resistant to inhibition of the PI3K-AKT-mTOR axis.

### 2.4 Imputing the effects of inhibitors based on a small measured dictionary

In our model breast cancer system, we were able to use PhEMD to assess the effects of a large panel of inhibitors on TGFβ-induced EMT. We found that these inhibitors could be grouped into clusters based on the similarity of their effects and embedded in low dimension (with an intrinsic dimensionality of three) to highlight complex, non-linear relationships between samples. Visualizing this embedding of inhibition conditions in 3D, we found that samples were distributed with varying density along a branched, continuous manifold. For example, the embedding space containing Cluster A inhibitors was characterized by high point density, while the embedding space containing Cluster D points was more sparsely populated (Figure 3c). We also noted that clusters often contained multiple inhibitors that targeted the same protein kinases. These findings suggested that we may have been able to capture the geometry of the drug-inhibition state space without measuring every single inhibition condition.

To test this hypothesis, we applied a previously published sampling technique to our PhEMD embedding [20]. The sampling technique used incompletely pivoted QR decomposition to identify “landmark points” (inhibition or control conditions) that approximately spanned the subspace of the single-cell sample embedding (Online Methods). Using this approach, we identified 30 landmark points that summarized our EMT perturbation state space (Supplementary Figure 8a). The 30 landmark points included samples from all 12 of Clusters A-L, suggesting they spanned all classes of experimental conditions in our experiment. To more fully assess whether the landmark points adequately captured the perturbation landscape of our full 300-sample experiment, we applied an accompanying out-of-sample extension technique to infer the embedding coordinates of all 300 samples relative to these 30 landmark points (Online Methods). The resulting embedding had a similar geometry to that of our original 300-sample PhEMD embedding, suggesting that the 30 landmark points were sufficient to capture the overall network structure of all 300 measured experimental conditions (Figure 3c, Supplementary Figure 8b). More generally, this finding supported the notion that redundancies may exist in a drug screen experiment, and that one may not need to measure an exhaustive set of perturbation conditions to uncover the underlying network geometry.

### 2.5 Validating the PhEMD embedding using external information on similarities between small-molecule inhibitors

We sought to validate our PhEMD drug-screen embedding by comparing the drug-drug similarities learned from our experiment (in the context of effects on EMT) to drug-drug similarities based on known drug-target binding specificities. To do so, we obtained drug-target binding specificity data from a recent study that used a chemical proteomic assay to identify all protein targets of each drug [21]. Since the previously published experiment and ours measured an overlapping set of inhibitors, they could be conceptualized as two complementary “views” of the same shared inhibitors. We conjectured that for the inhibitors shared between the two experiments, one view of the data might inform the other. Intuitively, this would support the notion that drugs with more similar protein targets action may tend to have more similar effects on EMT (and vice versa). Our approach to assessing this hypothesis was twofold: 1) We aimed to use a measure of inhibitor-inhibitor similarity, derived from the drug-target specificity data, to extend our PhEMD embedding and predict the effects of selected inhibitors on our model EMT system, and 2) We sought to use our PhEMD embedding to predict the drug-target specificity of inhibitors shared between the two drug-screen experiments.

#### 2.5.0.1 Predicting the effects of three selected inhibitors on breast cancer EMT relatively to the effects of measured inhibitors based on known drug-target binding specificities

For the first task, we sought to evaluate whether we could leverage known information on the mechanistic similarity between our inhibitors and additional inhibitors not measured in our experiment to predict the effects of these additional inhibitors on EMT. We selected saracatinib, ibrutinib, and dasatinib as three nonspecific Src inhibitors whose effects on EMT we wanted to predict. First, we generated a PhEMD embedding based on our CyTOF experimental results (not including the three selected inhibitors). Then, we obtained drug-target specificity data from a recently published inhibitor-profiling experiment [21] for inhibitors that overlapped between our experiment and the recently published one (including the 3 Src inhibitors of interest). We used the drug-target specificity data to compute pairwise cosine similarities between each of the 3 Src inhibitors and the samples in our initial PhEMD diffusion map embedding (that did not include the 3 inhibitors) (Online Methods). These pairwise similarities were used to perform Nystrom extension—a method of extending a diffusion map embedding to include new points based on partial affinity to existing points. In this way, we were able to predict the effects of the three Src inhibitors on breast cancer EMT relatively to inhibitors with known, measured effects (Online Methods).

To validate our extended embedding containing predicted Src inhibitor effects, we compared it to a “ground-truth” diffusion map embedding that used known (measured) CyTOF expression data for the 3 inhibitors and explicitly included the 3 inhibitors along with the rest in the initial embedding construction. Benchmarking our predictions against this ground-truth model, we found that our predictive model mapped the three inhibitors to the correct phenotypic space (Figure 4a-b). Specifically, saracatinib and ibrutinib were predicted to have an effect intermediate to those of specific MEK and EGFR inhibitors, and dasatinib was predicted to halt EMT less strongly than the other two Src inhibitors. These findings are consistent with ground-truth results based on direct CyTOF profiling and PhEMD-modeling of the three inhibitors (Figure 4b; Online Methods).

#### 2.5.0.2 Imputing the single-cell phenotypes of three unmeasured inhibitors based on drug-target similarity to measured inhibitors

We hypothesized that we could use drug-target information to not only relate unmeasured inhibitors to measured ones but also impute their single-cell compositions. To test this, we used the Nystrom-extended PhEMD embedding and dimensionality-reduced drug-target similarity data as inputs into a partial least squares regression model. We used this model to impute the cell subtype relative frequencies for the three unmeasured (imputed) Src inhibitors (Online Methods). As validation, we compared the predicted cell subtype relative frequencies to ground-truth CyTOF results (i.e., actual single-cell measurements) for the three inhibitors. PhEMD accurately predicted the cell sub-type relative frequencies for the three inhibitors compared to the null model (*P*=0.003, *P*=0.0008, *P*=0.009; Figure 4c-d).

To assess more generally whether PhEMD could be integrated with complementary data to accurately predict perturbation effects, we performed leave-out-out cross validation on all 39 inhibitors in our CyTOF experiment with known drug-target specificity data (Online Methods). We found that single-cell profile predictions leveraging PhEMD and knowledge of drug-target binding specificity were significantly more accurate than the null model (*P*=0.007). Altogether, these findings suggested that PhEMD offers information that can be integrated with additional data sources and data types to support not only comparison of samples directly measured but also prediction of single-cell phenotypes for additional, unmeasured samples.

#### 2.5.1 Predicting drug-target binding specificities based on PhEMD results from EMT perturbation experiment

We found that knowledge of drug-target binding specificity could be used to predict inhibitor effects in our model EMT system. We then sought to assess whether the reverse was true—whether the learned relationships between inhibitors from our EMT perturbation experiment could be used to predict drug-target binding specificities. If so, this would suggest that the two experiments measured two sets of inhibitor features that, while distinct between experiments, could be independently used to learn a consistent set of inhibitor-inhibitor relationships.

For this prediction task, we used the 39 inhibitors that were present in both the drug-target profiling experiment and ours, and that had at least 1 protein target identified by their experiment. We then computed leave-one-out predictions using the MAGIC imputation algorithm [22] and results from our EMT perturbation screen experiment to predict the drug-target binding specificities of each inhibitor (Online Methods). Prediction accuracy was defined as the correlation between predicted and measured drug-target binding specificities for a given drug. Our predictive model that incorporated PhEMD results into the prediction was significantly more accurate than the null model (*P*=6.57x10 ^−5^; Supplementary Figure 9a-b), suggesting that the inhibitor-inhibitor relationships learned from both experiments were consistent.

## 2.6 PhEMD highlights manifold structure of tumor samples in CyTOF and single-cell RNA sequencing experiments

To demonstrate an additional application of the PhEMD analytical approach, we used PhEMD to characterize the inter-sample heterogeneity in immune cell profiles of multiple tumor samples. We first applied PhEMD to a single-cell RNA-sequencing dataset consisting of the “healthy” (nontumor) cells of 17 melanoma biopsies [23]. The cell-state embedding identified a total of 11 cell subtypes with gene expression profiles consistent with previously reported subpopulations of B-cells, T-cells, epithelial cells, and macrophages (Figure 5a, Tirosh et al. [23]). When comparing patient samples, PhEMD identified the sample ‘Mel75’ as having a unique immune cell profile characterized by the greatest proportion of exhausted CD8^+^ cells. These cell-state and tumor-comparison findings corroborated previously published results on the immune cell subtypes and inter-sample heterogeneity present in this cohort. In addition to confirming prior findings, this analysis yielded an embedding that revealed the manifold structure of the single-cell sample state space. Samples at one end of the manifold were comprised mostly of B-cells and macrophages (G-3), samples at another end of the manifold consisted predominantly of CD8^+^ T-cells (G-2, G-4), and samples at the end of a third axis consisted of mostly macrophages (G-1). (Figure 5b, Supplementary Figure 10). While it is well-understood that a set of individual cells, such as those undergoing differentiation, may demonstrate manifold structure [24, 25], our PhEMD embedding suggested that a set of tumors from different patients with a shared phenotype (e.g., melanoma) may also lie on a continuous manifold.

To further explore this concept, we applied PhEMD to a mass cytometry dataset containing the T-cell infiltrates of 75 clear cell renal cell carcinoma (ccRCC) samples [26]. At the cellular level, our analysis recapitulated previous findings of important T-cell subpopulations present, including prominent CD8^+^ PD1^+^ CD38^+^ Tim-3^+^ exhausted T-cell (C-11) and CD4^+^ regulatory T-cell (C-2) populations (Figure 6a). To compare tumor samples to one another, we modeled the diversity in immune cell signatures as a tumor-sample embedding that could be visualized and partitioned to identify groups of similar samples (Figure 6b). As could be expected, the four healthy control samples all had similar immune cell profiles and were mapped close to one another on the tumor-sample manifold. This group of samples (G-2) demonstrated a mixed T-cell infiltrate comprised of both CD4^+^ and CD8^+^ T-cells. On the other hand, a different subgroup of tumor samples (G-3) was characterized by a marked predominance of exhausted CD8^+^ T-cells (C-11). In fact, progression toward one end of the tumor-space manifold represents a relative decrease in CD4+ T-cells (C-1, C-2) and relative increase in CD8+ PD1+ exhausted T cells (C-11) (Figure 6c, Supplementary Figure 11). This finding is supported by the initial report of substantial inter-patient variability in T-cell profiles especially related to CD8+ cells [26]. The detection of a subset of patients with exhausted T-cell enrichment may be of particular clinical interest, as immunotherapy agents that combat T-cell exhaustion have become a mainstay of advanced-stage ccRCC treatment but patients continue to have highly variable treatment responses [27, 28]. Future single-cell tumor-profiling experiments conducted to study treatment response may be able to use PhEMD as a tool to identify subgroups of patients that might especially benefit from PD-1 or PD-L1 inhibitor immunotherapy.

## 3. Discussion

Herein, we have shown that we can successfully map the space of experimental conditions, measured as a set of single-cell samples, using our proposed PhEMD embedding technique. Each experimental condition embedded represents a setting of an experimental variable, such as a unique drug with which a cell population is stimulated. We extensively studied the Py2T murine breast cancer cell line treated TGF-β and perturbed with over 200 kinase inhibitors, measured using mass cytometry. In this experiment, PhEMD revealed the structure of the kinase inhibitor space based on each drug’s effect on the Py2T cell populations undergoing EMT. This network of inhibitors was found to have low-dimensional structure, with drugs mapping to one of three main axes. We have shown that the embedding produced by PhEMD is useful in several ways:

1. Visualization and intuitive understanding of the experimental variable (i.e., single-cell sample) state space.
2. Clustering and extraction of similar settings of experimental variables (e.g., similar drugs with respect to their measured effects on a given cell population).
3. Characterization of clusters and axes of variability in the experimental variable state space in terms of biologically-interpretable differences in the types and abundances of cell subpopulations present in each sample.
4. Extension of the experimental variable state space through inference of unmeasured experimental settings based on similarity to existing (measured) settings.

Most notably, PhEMD can enable a new paradigm of searching for effective therapeutic agents (e.g., drugs that perturb a cancer cell population) by identifying a small dictionary of prototypical drugs that collectively capture the network geometry of a larger drug set. We demonstrated this application by computing a dictionary of 30 experimental conditions and showing that these 30 kinase-inhibitor and control conditions were sufficient to capture the network geometry of the 300-sample state space. More concretely, we showed that the true 300-sample network geometry could be recovered by relating all 300 inhibition and control conditions to the 30 dictionary samples. This finding has the potential to reduce experimental burden in future drug discovery efforts. For example, one can first apply PhEMD to measurements obtained using one profiling technique (e.g., mass cytometry) to compute a small set dictionary samples from a large set of candidates. One may then further investigate this small set of dictionary samples using complementary technologies that may be more limited in scale (e.g., single-cell RNA sequencing). This approach can be used to facilitate a systematic nomination and feasible investigation of the most promising therapeutic agents.

We validated our drug-screen PhEMD embedding by comparing it to recently-published drugtarget specificity data on the same inhibitors. We first showed that by using a measure of inhibitor-inhibitor similarity derived from drug-target binding specificity data, we could predict the effects of unmeasured inhibitors on our model EMT system. We then showed that the reverse was also true: that by using our PhEMD embedding, we could predict drug-target specificity with better accuracy than a null model. Our observation that the drug-target specificity data could be used to inform a prediction of inhibitor effect in our model system and vice versa was consistent with the notion that the two experiments were two views of the same data points (i.e., inhibitors). Both experiments learned intrinsic properties of the inhibitors that could be used to derive similar yet distinct insights into relationships between inhibitors.

In addition to validating the learned relations between inhibition conditions, our assessment of PhEMD as a potential predictive tool highlighted the ability of PhEMD results to be integrated with additional data sources and data types for even larger and richer analyses. By using drug-target specificity data from another large-scale inhibitor profiling experiment, we could leverage a powerful, out-of-sample extension method known as *Nystrom extension* to insert additional inhibitors into the embedding. Thus, we were able to accurately predict the effects of inhibitors not directly measured in our experiment on TGF-β-induced breast cancer EMT. This approach may be useful for analyzing drug-screen experiments, as it enables a mapping of a small set of drugs (e.g., dictionary points) measured at single-cell resolution to be extended to include additional drugs. Moreover, we believe this application is not limited to analyzing perturbation screens and can be useful for imputing the phenotypes of samples (of any type) that are not directly included in a single-cell sequencing experiment. For example, examining a cohort of patients in which only some patients were biopsied and genomically profiled, one could potentially incorporate a non-genomic based measure of patient–patient similarity (e.g. based on clinicopathologic features) to predict the single cell-based phenotypes of all patients in the cohort.

We explored the applicability of PhEMD to other experimental designs besides drug screens by applying PhEMD to three additional datasets. These analyses revealed that PhEMD reliably uncovered manifold structure in the single-cell sample space that was biologically interpretable and reasonable based on the observed proportions of the samples’ cell subpopulations. As could be expected, in our simulated dataset, the PhEMD sample embedding modeled a trajectory that consisted of the samples in which cell density was gradually modulated along one axis of the underlying cell state tree. When applying PhEMD to a melanoma dataset consisting of immune cell measurements from tumor biopsies, we found that PhEMD revealed “trajectories” of patients, with the most notable axis consisting of patients with an increasing proportion of CD8^+^ T-cells. PhEMD applied to a dataset of tumor-infiltrating T-cells in renal cell carcinomas similarly revealed a prominent trajectory of patients with increasing exhausted CD8^+^ T-cells. By organizing patients and their cell populations in this way, PhEMD highlighted important biological differences between patients. It is possible that the abundance of tumor-infiltrating, exhausted T-cells may predict response to immunotherapy, although additional studies are needed to assess this. If true, our findings and the PhEMD method may be useful in paving the way for developing personalized cancer treatment regimens involving immunotherapy.

Through our analyses, we demonstrated that PhEMD can be used to characterize mass cytometry and single-cell RNA-sequencing data, though we believe PhEMD may be applied to data generated by other single-cell profiling platforms as well. PhEMD is generalizable not only to data type but also experimental setup. We can envision many experiments that may benefit from PhEMD— for example, comparisons of samples pre- and post-treatment (or receiving different treatments), time-series analyses of cells undergoing transition processes, and organization of heterogeneous-yet-related samples for the purpose of disease subtyping. Additionally, applying PhEMD to large-scale functional genomics (e.g., single-cell CRISPR) screens may yield sample embeddings that highlight complex relationships between genes. We have demonstrated in our analysis of over 1.7 million cells across 300 samples and five experimental runs that PhEMD is highly scalable and robust to batch effect. As single-cell datasets of increasingly large sample size are generated, we believe PhEMD offers the efficiency, flexibility, and model interpretability necessary to fully leverage the information that single-cell data offers.

### 3.1 Available Code & Data

PhEMD (“Phenotypic Earth Mover’s Distance”) takes as input a list of *N* matrices representing *N* single-cell samples. An R implementation of PhEMD can be installed from https://github.com/wschen/phemd, and we plan to make the package available on Bioconductor soon (package: ‘phemd’).

## 4. Author Contributions

W.C., N.Z., G.W., B.B., and S.K. conceived of the study. W.C. and D.v.D. implemented the PhEMD algorithm and performed all computational analyses. N.Z. performed all single-cell profiling experiments and data quality assessments. W.C., N.Z., B.B., and S.K. interpreted the results and wrote the manuscript.

## 5. Online Methods

### 5.1 The PhEMD analytical approach

In single-cell data, each cell is characterized by a set of features, such as protein or transcript expression levels of genes. The purpose of measuring these expression-based features for each cell (e.g., via single-cell RNA-seq or mass cytometry) is to answer biological questions especially related to the cell subpopulations present in a sample. In particular, the features may be used for defining phenotypes of cells [29, 30], resolving cellular dynamics using transition-process modeling [31–33], and studying signaling networks [34, 35]. In sum, the features are shared, quantitative characteristics of cells that may be used to organize a set of cells into a data geometry. An analogy can be made when attempting to compare single-cell samples rather than individual cells. A sample is a collection of cells. In order to compare single-cell samples for the purpose of organizing a set of cell collections (e.g., different patient samples or perturbation conditions), one must first determine useful features for a cell collection. Previous studies have shown that cell subtypes are highly useful features that are shared across all samples and can be quantitatively measured. Moreover, they can be used to represent single-cell samples efficiently for downstream analyses (we expand on this notion in Supplementary Note 2). Just as transcript counts can be measured for selected genes in a single cell, so can cell counts be measured for selected cell subtypes in a cell collection.

We use Monocle 2 for the task of defining cell subtypes [33]. Monocle 2 performs reversed graph embedding on high-dimensional single-cell data to both identify unique cell subpopulations and relate them to one another on a manifold (i.e., “tree”). By applying Monocle 2 to an aggregate of cells in a single-cell experiment, we can represent a biological sample as the relative frequency of cells in each cell subtype. This representation of single-cell samples is consistent with the “signatures-and-weights” representation of multidimensional distributions, first formalized by Rubner et al. [36], that was found to yield optimal data representation efficiency in other computer vision applications (Supplementary Note 2). In our case, a “signature” can be thought of as a distinct cell subtype (e.g., memory B-cells or CD8+ effector T-cells), and the corresponding “weight” represents the proportion of cells in a given sample assigned to the cell subtype. However, comparing single-cell samples represented as such is still a non-trivial task (Supplementary Note 3). Many studies represent single-cell samples as their cell subtype composition and use known class labels (e.g., normal lung vs. lung adenocarcinoma) to group samples and perform class-based comparisons (e.g., identifying cell subtypes enriched in a disease state) [37, 38]. However, this approach is limited to comparing a few predefined classes of samples and does not reveal insights into intra-class heterogeneity. Other studies organize a set of many single-cell samples based on their relative frequency of one or a few important cell subtypes [30,39,40]. However, this approach requires *a priori* knowledge of the most important cell subtypes and does not provide a complete view of sample-to-sample dissimilarity, especially in the context of high intra-sample cellular heterogeneity.

We posit that the ideal metric for comparing samples should take into account both the difference in weights of matching bins (e.g., number of CD8+ T-cells) for all bins and the dissimilarity of the bins themselves (e.g., intrinsic dissimilarity between CD8^+^ and CD4^+^ T-cells). Earth mover’s distance (EMD) is a nonparametric metric that can capture both of these concepts to yield a final singular measure of distance, or dissimilarity, between two samples [36]. EMD can be conceptualized as the minimal amount of “effort” needed to move mass (e.g., cells) between bins of one histogram so that its shape matches that of the other histogram (i.e., all matching bins of two histograms have the same counts). Mathematically, EMD is defined by the following optimization problem:

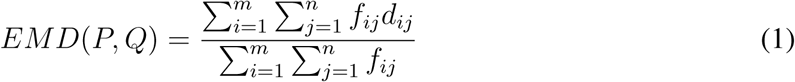

Such that 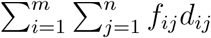 is minimized subject to the following constraints:

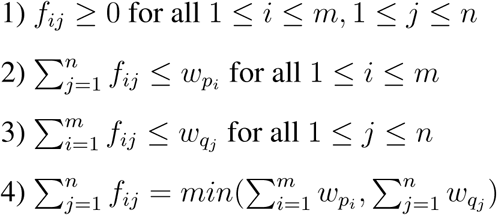

#### Definition 1

*Earth Mover’s Distance as an optimization problem. P* = (*p*_1_, *wp*_1_), *…*, (*p*_*m*_, *wp*_*m*_), *where p*_*i*_ *represents histogram bin i in the initial starting signature P and represents the amount of “mass” present in the bin. Similarly, Q* = (*q*1, *wq*_1_), *…*, (*qn, wq*_*n*_), *where q*_*j*_ *represents histogram bin j in the final signature Q and represents the amount of “mass” present in the bin. 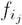 represents the “flow” of mass from bin p*_*i*_ *to bin q*_*j*_*. 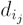 represents the “ground distance” between bins p*_*i*_ *and q*_*j*_*. Constraint 1 ensures that P and Q are the starting and final signatures respectively. Constraints 2 and 3 ensure that no more mass is moved from any bin pi than is present initially. Constraint 4 ensures that the maximum amount of mass is moved as possible, i.e., the final signature most closely resembles Q as possible.*

EMD has been used in various applications including image retrieval [36, 41], visual tracking [42], and melodic similarity musical analysis [43]—all tasks that require accurate comparison of multidimensional distributions (analogous to comparing single-cell samples). Additionally, a prior study demonstrated proof-of-concept that Earth Mover’s Distance can be used effectively to differentiate flow cytometry samples of phenotypically distinct individuals [44]. By design, EMD is a distance measure between probability distributions that is particularly invariant to small shifts in data (i.e., noise or technical variability) across samples [36, 44]. EMD also gives a “complete” measure of overall dissimilarity between two samples, largely attributable to the fact that it takes into account both the difference in height of corresponding histogram bins between samples (e.g., number of CD8+ cells) and the concept that certain bins (e.g., cell subtypes) have a smaller “ground distance” (i.e., are more similar) than others. Including ground distance between bins in the EMD computation allows us to incorporate the idea that it requires more “effort” to move mass to a faraway bin than to a nearby bin (i.e., it requires more effort to convert cells to a more dissimilar cell signature than to a more similar cell signature). In our application, we define the ground distance between two cell subtypes as the manifold distance (“tree-distance”) between the cluster centroids of the two cell subpopulations representing the subtypes (Supplementary Note 4, Figure 1c-d, Suppl. Fig. 2).

Leveraging these features of EMD, we developed PhEMD as a novel approach to comparing single-cell samples (Figure 1b). PhEMD first aggregates cells from all biological samples and applies Monocle 2 to model the cell-state space. Monocle 2 simultaneously identifies all cell sub-types and relates them in a tree-based embedding. After constructing the Monocle 2 embedding of the cell-state space, PhEMD represents each biological sample to be compared as a frequency histogram capturing relative abundance of each cell subtype. PhEMD then uses EMD, incorporating tree-distance as ground-distance between bins, to compare two relative abundance histograms and derive a single value representing the dissimilarity between two single-cell samples. PhEMD computes EMD pairwise for each pair of samples to generate a distance matrix representing sample–sample dissimilarity. This distance matrix can be directly inspected to investigate pairwise sample-sample dissimilarity and embedded in low-dimensional space using diffusion maps to view sample–sample relationships in the context of overall network structure (Supplementary Note 6) [45]. Diffusion maps are useful in this case as they learn a nonlinear mapping of samples from high-to low-dimensional space that captures both local and global structure and has intrinsic denoising properties. PhEMD constructs a diffusion map embedding and clusters embedded samples to identify groups of similar samples based on the compositional similarity of their respective cell populations.

### 5.2 Py2T cell culture and stimulation

Py2T cells were obtained from the laboratory of Gerhard Christofori, University of Basel, Switzerland [46]. Cells were tested for mycoplasma contamination upon arrival and regularly during culturing and before being used for experiments. Cells were cultured at 37°C in DMEM (Sigma Aldrich), supplemented with 10% FBS, 2 mM L-glutamine, 100 U/mL penicillin, and 100 μg/mL streptomycin, at 5% CO2. For cell passaging, cells were incubated with TrypLE™ Select 10X (Life Technologies) in PBS in a 1:5 ratio (v/v) for 10 minutes at 37°C.

Human recombinant TGF-β_1_ was purchased from Cell Signaling Technologies as lyophilized powder and was reconstituted in PBS containing 0.1% carrier protein, according to the manufacturer’s protocol to 400 ng/mL. The stock solution was kept at -20°C until use. For daily treatment, TGF-β_1_ stock was diluted into medium to 40 ng/mL working concentration. Following small-molecule inhibitor treatment, 10 μL TGF-β_1_ was added to the cells for a final concentration of 4 ng/mL. As a control, PBS containing carrier protein diluted with growth medium was used.

### 5.3 Small molecule inhibitors

A library of 233 small molecule kinase inhibitors was purchased from Selleckchem (Supplementary Table 1). Small-molecule inhibitors were distributed within the 60 inner wells of five separate 96-well format deep well blocks with exception of wells within row E, which contained DMSO. Stock solutions of 2 mM small molecule inhibitor in DMSO were kept at -80°C until used. For daily treatment, the stock solution was equilibrated at room temperature for 1 h and then 5 μL of stock solution was added 995 μL of medium. Small-molecule inhibitor (or DMSO) was added to cells once per day, immediately after the cell growth media change and before application of TGF-β_1_. Small-molecule inhibitor treatment was performed by adding 10 μL of pre-diluted reagent to the cells in 80 μL cell growth medium; this resulted in a final concentration of 1 μM of small-molecule inhibitor and 0.1% DMSO.

### 5.4 Chronic kinase inhibition screen

For the chronic inhibition experiment, Py2T cells were seeded in 96-well plates (TPP, Techno Plastic Products AG) with a seeding density of 1800 cells per well in 80 μL of growth cell media. Only the 60 inner wells were used for analysis. In order to acquire sufficient sample size, five 96-well plates were used for single condition. After seeding, cells were allowed to recover for 36 h to reach 50% confluence. Cells were treated simultaneously with TGF-β_1_ or vehicle (PBS) and small-molecule inhibitor or vehicle (DMSO) for 5 days, and medium was changed daily. All pipetting procedures were performed at room temperature using a Biomek FX Laboratory Automation Workstation (Beckman Coulter) supplied with 96-well pipetting pod.

### 5.5 Cell harvesting

The cell harvesting protocol was performed using a Biomek® FX Laboratory Automation Workstation. The cell growth medium was removed using the multiple aspiration pipetting technique, and cells were washed twice with 37°C PBS. Dissociation reagent TrypLE™ Select 10X (Life Technologies) was diluted into PBS at a 1:5 ratio (v/v) was added to the cells and incubated for 10 min at 37°C. Cells were detached from plates. Five identically treated 96-well plates were combined into a single deep well block and were fixed for 10 min with PFA at the final concentration of 1.6% v/v. PFA was blocked with the addition of 600 μL 10% BSA in CSM. The cells were centrifuged for 5 min at 1040 x g, 4°C. The supernatant was removed and the cells were resuspended in 300 μL of -20°C MeOH. Samples were then transferred onto dry ice and to -80°C storage.

### 5.6 Metal-labeled antibodies

Antibodies were obtained in carrier/protein free buffer and labeled with isotopically pure metals (Trace Sciences) using MaxPAR antibody conjugation kit (Fluidigm) according to the manufacturer’s standard protocol. After determining the percent yield by measurement of absorbance at 280 nm, the metal-labeled antibodies were diluted in Candor PBS Antibody Stabilization solution (Candor Bioscience GmbH) for long-term storage at 4°C. Antibodies used in this study are listed in Supplementary Table 2.

### 5.7 Mass-tag cellular barcoding and antibody staining

Cell samples in methanol were washed three times with Cell Staining Media (CSM, PBS with 0.5% BSA, 0.02% NaN3) and once with PBS at 4°C. The cells were then resuspended at 1 million cells/mL in PBS containing barcoding reagents (^102^Pd, ^104^Pd, ^105^Pd, ^106^Pd, ^108^Pd, and ^110^Pd; Fluidigm) were conjugated to bromoacetamidobenzyl-EDTA (BABE, Dojindo) and two indium isotopes (^113^In and ^115^In, Fluidigm) were conjugated to 1,4,7,10-tetraazacy-clododecane-1,4,7-tris-acetic acid 10-maleimide ethylacetamide (mDOTA, Mycrocyclics) following standard procedures [47, 48]. Cells and barcoding reagent were incubated for 30 min at room temperature. Barcoded cells were then washed three times with CSM, pooled and stained with the metal-conjugated antibody mix (Supplementary Table 2) at room temperature for 1 h. Unbound antibodies were removed by washing cells three times with CSM and once with PBS. For cellular DNA staining, an iridium-containing intercalator (Fluidigm) was diluted to 250 nM in PBS containing 1.6% PFA, added to the cells at 4°C, and incubated overnight. Before measurement, the intercalator solution was removed and cells were washed with CSM, PBS, and doubly distilled H_2_O. After the last wash step, cells were resuspended in MilliQ H_2_O to 1 million cells/mL and filtered through a 40-μm strainer.

### 5.8 Mass cytometry data processing

EQ Four Element Calibration Beads (Fluidigm) were added to the cell suspension in a 1:10 ratio (v/v). Samples were measured on a CyTOF1 system (DVS Sciences). The manufacturer’s standard operation procedures were used for acquisition at a cell rate of ∼300 cells per second as described previously [49]. After the acquisition, all FCS files from the same barcoded sample were concatenated using the Cytobank concatenation tool.

Data were then normalized [50] and bead events were removed. Cell doublet removal and de-barcoding of cells into their corresponding wells was done using a doublet-free filtering scheme and single-cell deconvolution algorithm [47]. Subsequently, data were processed using Cytobank (http://www.cytobank.org/). Additional gating on the DNA channels (^191^Ir and ^193^Ir) was used to remove remaining doublets, debris, and contaminating particles. Final events of interest were exported as .csv files.

### 5.9 In-depth analysis of breast cancer EMT cell-state space and drug-inhibitor manifold from a single mass cytometry run

CyTOF measurements of cells undergoing unperturbed and perturbed EMT were generated and processed as described above. Data were then pooled from all experimental conditions, taking an equal random subsample from each condition to generate the cell state embedding. Cell state definitions and relationships were modeled with Monocle 2, using the ‘gaussianff’ expression model and sigma (noise) parameter of 0.06. Subsequently, all cells from all experimental conditions were assigned a cell subtype using a nearest-neighbor approach (Supplementary Note 7). Deconvolution was then performed to generate relative frequency histograms representing distribution of cells across all cell subtypes for each inhibition condition. EMD was computed pairwise between single-cell samples, using manifold distance (i.e., Monocle 2 pseudotime) as a measure of intrinsic dissimilarity between cell subtypes for the EMD ground-distance matrix (Fig. 2, Suppl. Fig. 1). The resulting sample-to-sample distance matrix was embedded using the ‘destiny’ Bioconductor `R` package [51] and partitioned using hierarchical clustering to highlight inhibitors with significant effects on EMT or similar effects to one another.

### 5.10 Intrinsic dimensionality analysis of the EMT perturbation state space

To assess the intrinsic dimensionality of the EMT perturbation state space, we applied the bias-corrected maximum likelihood estimator approach [52]. We computed the sample-to-sample distance matrix for the 60 samples measured in a single CyTOF run as described above and estimated intrinsic dimensionality of this embedding using the ‘ider’ `R` package [53]. Intrinsic dimensionality was estimated over a range of values for knn parameter *k* from 1 through 20. The final value of intrinsic dimensionality was determined by examining the stable estimated value across a range of sufficiently large values for *k*.

### 5.11 Integrating batch effect correction to compare 300 EMT inhibition and control conditions measured in 5 experimental runs

CyTOF measurements of cells undergoing unperturbed and perturbed EMT were generated and processed as described in the above sections. Given slight differences in CyTOF marker panels between batches, only markers shared across all batches (n = 31) were used for downstream analyses. Data were pooled from all experimental conditions on a per-batch basis. Expression values were then linearly scaled for each gene to ensure all values were positive and in the same range across batches. After this initial normalization, an equal random subsample of cells from each batch (20,000 x 5) was used as the input for canonical correlation analysis (CCA) [18]. CCA mapped expression data from each batch into an aligned, 12-dimensional space shared by all batches. Cell state definitions were modeled using the Louvain community detection method included in Seurat (’FindClusters’), using the 12 dimensions of the CCA-aligned space as input and specifying a clustering resolution of 0.7. The cell state space was visualized by applying the PHATE dimensionality reduction method [7] on the same input data as was used for Louvain community detection.

All cells from all experimental conditions were assigned a cell subtype using a nearest-neighbor approach (Supplementary Note 8). Deconvolution was then performed to determine the cell subtype distribution of each inhibition condition. Using this cell subtype-based representation of inhibition conditions, EMD was computed pairwise between single-cell samples. Since Monocle 2 was not compatible with CCA-aligned data, we defined the ground distance (i.e. intrinsic dissimilarity) between cell subtypes as the Euclidean distance between their respective centroids in the aligned, 12-dimensional space. The resulting sample-to-sample distance matrix was embedded using the ‘destiny’ Bioconductor R package [51] and partitioned using hierarchical clustering to identify 12 clusters of inhibitors with similar effects on EMT. The intrinsic dimensionality (i.d.) of this multi-batch, 300-sample PhEMD embedding was estimated as described in the above section, taking into account i.d. estimations over a range of *k* values from 1 through 100.

### 5.12 Imputing the effects of inhibitions based on a small measured dictionary

To assess whether the network geometry of all 300 inhibition and control conditions could be captured using a smaller subset of conditions, we applied a previously published sampling technique for identifying landmark points of an embedding [20]. First, the PhEMD distance matrix containing pairwise distances between our 300 experimental conditions was converted to an affinity matrix using a Gaussian kernel (*σ* = 2) and Markov-normalized to obtain probabilities. The incompletely pivoted QR-based (ICPQR) dimensionality reduction technique was then applied, using a *μ* distortion parameter of 0.01, to identify 30 landmark points. The landmark points identified were then used to impute the geometric coordinates of the remaining (non-landmark) points using the out-of-sample extension technique associated with ICPQR [20]. The result was an 30-dimensional embedding of all 300 samples. We computed a 300x300 distance matrix based on the pairwise Euclidean distances between samples in this 30-dimensional space and then embedded using the ‘destiny’ Bioconductor `R` package [51].

### 5.13 Incorporating drug-target binding specificity data to extend the PhEMD embedding and predict the effects of unmeasured inhibitors on TGF**β**-induced breast cancer EMT

We hypothesized that we could predict the influence of additional inhibitors on TGFβ-induced EMT based on knowledge of inhibitor-inhibitor similarity from another data source. To test this, we obtained drug-target specificity data from a previously published experiment [21] for a set of 39 inhibitors that overlapped between our experiment and theirs. We then selected saracatinib, ibrutinib, and dasatinib as three nonspecific Src inhibitors whose drug-target specificity data were known and whose effects on EMT we wanted to predict. Next, we generated a PhEMD embedding based on our CyTOF experimental results (not including the three selected inhibitors). To predict the effects of the three inhibitors on EMT relatively to other inhibitors in our experiment, we performed Nystrom extension on the diffusion map embedding. All 39 inhibitors that were found to have an effect on EMT in our experiment and that had known drug-target specificity profiles were included in the Nystrom extension. Pairwise distances between each “extended” point and each existing point in the original diffusion map were required for Nystrom extension. These distances were based on the similarity of drug-target specificity profiles between the two inhibitors, defined as (1*-cosine similarity*)^20^ *4 for all pairs of inhibitors with known drug-target specificity profiles. The remaining pairwise distances were imputed based on known PhEMD-based inhibitor-inhibitor dissimilarity and known pairwise drug target specificity-based dissimilarity using the MAGIC imputation algorithm [22].

We observed a global shift in embedding coordinates between the original diffusion map (based on PhEMD distances) and the Nystrom extension points (based on normalized cosine similarity using drug-target specificity data). This was likely due to a difference in scale between PhEMD-based distances and cosine similarity-based distances. Nonetheless, we were able to use the Nystrom extension points alone to predict the effect of the three selected inhibitors on EMT. First, we visualized the Nystrom extension embedding to show the predicted relation of the three inhibitors to other inhibitors with known (measured) effects on EMT. Next, we used partial least squares regression (‘pls’ `R` package) to predict the cell subtype relative frequencies that would result from applying the inhibitors to breast cancer cells undergoing TGFβ-induced EMT. Input variables for the regression model were the Nystrom extension embedding coordinates and first 5 principal components of the drug-target-based inhibitor similarity matrix transformed using principal components analysis (PCA). To validate our findings, we measured the three selected inhibitors directly using CyTOF and included them along with the rest of the inhibitors in the PhEMD analysis pipeline. We compared the actual to the predicted cell subtype relative frequencies and the actual to the predicted embedding coordinates relative to other similar, “nearby” inhibitors. To assess prediction accuracy, we compared our prediction error to the prediction error of the null hypothesis modeled by first randomizing the PhEMD-based and drug target specificity-based distance matrices and then generating a predictive model in the same way as in the alternative model. Prediction error was defined as the EMD between the predicted and actual (measured) cell subtype relative frequency distributions. The null hypothesis was modeled as a distribution of EMDs generated by randomizing the PhEMD-based and drug target specificity-based distance matrices 100 times and subsequently imputing cell subtype frequencies. P-values were computed by computing the z-score of the prediction error based on the null distribution and applying a one-sided test at a significance level of 0.05.

To more comprehensively assess PhEMD as a predictive tool, we performed leave-one-out cross validation on the 39 inhibitors with known (measured) cell subtype relative frequencies and drug-target specificity data. For each inhibitor, we constructed a PhEMD embedding based on known measurements of the 39 others and performed Nystrom extension to impute the relationship between the inhibitor and the measured ones. We then constructed a partial least squares regression model using the same input variables as above to predict the cell subtype relative frequencies of the inhibitor. Prediction error was defined the same as above (i.e. EMD between predicted and actual cell subtype relative frequency distributions). The null model was also defined in the same way as above by randomizing the PhEMD and distance matrices 100 times for the prediction of each inhibitor. To determine whether our alternative model was effective, we assessed whether the prediction errors in the alternative model (n=40) were collectively lower than the EMDs in the null model (n=4,000) using the Kolmogorov-Smirnov test.

### 5.14 Predicting drug-target binding specificities based on PhEMD results from EMT perturbation experiment

We hypothesized that if the PhEMD embedding were meaningful, it would have predictive power. In order to test this, we used the PhEMD embedding of inhibitors to predict the inhibitors’ drug-target binding specificities. The drug-target binding specificity data were obtained from a previously published study that used a chemical proteomic approach to identify the protein targets of many clinical kinase inhibitors [21]. We chose to predict the profiles of 39 inhibitors that were present in both the drug-target binding specificity experiment and ours, and that had at least 1 protein target identified by the binding specificity experiment. Next, we computed a 39-by-39 knn kernel (k=3) using the PhEMD inhibitor-inhibitor distances and then row-normalized the resulting matrix to 1 to turn it into a Markov operator. We then performed a leave-one-out cross validation, in which we set one of the inhibitor target values (i.e., drug-target binding specificity profiles) in the Klaeger et al. data to be unknown. Note that a drug-target binding specificity profile was represented as a vector of length 270, which represented the binding specificity between the drug and each of 270 potential protein targets. We predicted the drug-target binding specificity values using the MAGIC imputation method with the PhEMD Markov operator as input and a diffusion parameter *t* of 2. We computed leave-one-out predictions for each of the 39 inhibitors. To quantify the performance of our predictive model, we computed Pearson correlation between the original ground-truth (experimentally measured) target values and the predicted values. To determine the accuracy of our predictions, we compared our results to a null model, in which we randomized the PhEMD matrix 1,000 times and each time ran the prediction using this randomized matrix. Prediction accuracy (Pearson correlations) of our alternative model was compared to that of the null model using a one-sided Kolmogorov-Smirnov test.

### 5.15 Generation and analysis of dataset with known ground-truth branching structure

To evaluate the accuracy of the PhEMD analytical approach, high-dimensional single-cell data were generated based on a modified version of the “artificial tree” test case used in a prior study [7]. In the basic tree structure represented in Figure 2a, the first branch (C-4) consisted of 100 cells whose expression values increased linearly in the first four dimensions and were zero in all other dimensions. The branch then bifurcated into two branches, C-5 and C-3, which consisted of cells with constant expression in the first four dimensions (value equal to the endpoint of branch C-4) and which increased linearly in dimensions 5-8 and 9-12 respectively. Expression of all other features for cells in these branches was zero. The remaining branches were constructed similarly. Thirty additional cells were added to the endpoints of each branch. Six non-informative features were added (that consist exclusively of zeros) for a total of fifty dimensions. Zero-centered Gaussian noise was added to all expression values. To simulate a unique biological sample, the basic tree is constructed as described above and a total of 900 additional, linearly-spaced cells were distributed across one or multiple selected branches. Finally, all data were z-score normalized by feature.

We applied PhEMD to the z-score normalized data as outlined in Figure 1. First, the tree structure was modeled by Monocle 2 based on cells aggregated from all biological samples. Then, the relative frequency of cells across different cell subtypes was computed for each sample. EMD was computed pairwise for all cells using Monocle 2 “tree-distance” as a measure of ground-distance between cell subtypes and a final embedding of biological samples was generated using the ‘destiny’ Bioconductor `R` package (Fig. 3).

### 5.16 Analysis of melanoma single-cell RNA sequencing dataset

Data from a prior single-cell RNA-sequencing experiment were downloaded from the NCBI Gene Expression Omnibus website, accession number GSE72056 [23]. These data contained read-count expression values that were log TPM-normalized values. 2 of the 19 samples were excluded from analysis due to low cell yield of immune cells. Initial feature selection was performed by selecting 44 features found in the initial publication characterization of this dataset to distinguish between key cell types [23]. The Monocle 2 tree-based model of the cell-state space was constructed using the ‘gaussianff’ expression model recommended by the Monocle 2 tutorial with a sigma (noise) parameter of 0.02. The remaining PhEMD analysis pipeline was completed as described in ‘Characterizing effects of chronic drug-inhibitor application on breast cancer cells undergoing EMT’; a final embedding of biopsy samples was generated using the ‘destiny’ Bioconductor `R` package and partitioned using hierarchical clustering.

### 5.17 Analysis of clear cell renal cell carcinoma dataset

CyTOF data from a recent publication characterizing the immune landscape of clear cell renal cell carcinoma were downloaded from https://premium.cytobank.org/cytobank/projects/875 [26]. Cell data were filtered and normalized using the method described in Online Methods section ‘Mass cytometry data processing’. The Monocle 2 tree-based model of the cell-state space was constructed using the ‘gaussianff’ expression model with a sigma (noise) parameter of 0.02. The remaining PhEMD analysis pipeline was completed as described in ‘In-depth analysis of breast cancer EMT cell-state space and drug-inhibitor manifold from a single CyTOF experiment’.

## 8. Supplementary notes

### Supplementary Note 1: Leveraging single-cell resolution to distinguish samples that are indistinguishable by bulk expression analysis

Bulk expression analysis may reveal trends that inadequately reflect the true differences between biological samples. For example, a prior report studying pulsatile expression of p53 in cells before and after γ-irradiation treatment found that on bulk analysis, the average amplitude of pulses (i.e., magnitude of response to treatment) was greater with increasing dose of irradiation [1]. A natural conclusion from this observation may be that cells express increased p53 in response to irradiation-induced DNA damage. However, the group then performed the same experiment but obtained single-cell instead of bulk measurements. This experiment revealed that the pulse amplitude for a given cell was actually constant and independent of irradiation dose; the change in average pulse amplitude on bulk analysis was attributable to changing proportions (i.e., preferential survival and/or proliferation) of certain cell subpopulations rather than changes in individual cells themselves. Without single-cell resolution, this distinction could not be made.

In addition to lacking the resolution to explain observed trends, bulk measurements may fail to detect true biological differences between experimental conditions altogether. To demonstrate this concept more concretely and highlight the utility of single-cell analytical approaches for distinguishing between biological samples, we computationally modeled a multi-sample dataset consisting of immune cells with collectively variable expression of CD4 and CD8 (Suppl. Fig. 1a). Each sample was a cell population that fit one of four distribution patterns. Group A samples each consisted of a homogeneous immune cell population characterized by intermediate expression of both CD4 and CD8. Group B samples each consisted of two similarly-sized immune cell subpopulations: one CD4^+^ and one CD8^+^ subpopulation. Group C samples consisted of a mixture of CD4^+^, CD8^+^, and CD4/CD8 double-positive (DP) immune cells. Group D samples consisted of one CD4^+^ and one CD8^+^ subpopulation of roughly equal size and one additional rare subpopulation of CD4/CD8 double-negative (DN) immune cells. Note that these immune cell subtypes (CD4^+^, CD8^+/^, DP, and DN) have been reported to exist in normal thymus as well as various disease states (e.g., breast and hematologic malignancies [2, 3]). Our simulated experiment consisted of 32 samples in total (8 of each of Groups A-D). By design, the bulk (average) expression of CD4 and CD8 for each sample was roughly the same for all samples, regardless of differences in cell subpopulation characteristics.

Our goal was to relate the 32 samples to one another in a biologically meaningful way. This could be done by generating a low-dimensional embedding that could be visualized to view the similarity of any two samples relative to the rest and identify groups of similar samples. We first attempted to do so using bulk measurements. We generated a sample-sample distance matrix by computing pairwise (Euclidean) distances between each pair of samples, with each sample represented as its average protein (i.e., CD4 and CD8) expression. We then embedded this distance matrix using a diffusion map. The result was an embedding that failed to differentiate samples based on biologically important differences. Specifically, samples of the same known, groundtruth subtype (i.e., Group A-D) failed to map to similar parts of the resulting embedding (Suppl. Fig. 1b).

A better approach to comparing these samples was to compare the presence and abundance of all single-cell subpopulations in each sample. We formalize an approach (”PhEMD”) in this manuscript and demonstrate that it can be used to effectively distinguish single-cell samples from one another that cannot be distinguished based on bulk or average expression patterns. In this particular example, the PhEMD diffusion map embedding vastly outperformed the bulk approach described above and successfully differentiated samples based on biologically important differences in cell subpopulation characteristics and proportions (Suppl. Fig. 1c).

### Supplementary Note 2: Motivation for “cell subtype” features

Individual cells can be thought of as points in an *n*-dimensional space, in which each dimension represents the expression level of a specific gene or protein marker. Thus, a single-cell sample containing thousands of cells can be thought of as a distribution of points in an *n*-dimensional space. Our goal is to perform pairwise comparisons of samples. Naturally, it follows that we aim to compare the *n*-dimensional distributions by computing a distance between two distributions. A naive geometric approach is to bin the distribution into a n-dimensional joint histogram with b bins in each dimension. However, this results in *b*^*n*^ bins, which implies extreme sparsity and computational intractability, as *n* is generally in the tens to thousands for single-cell genomic datasets.

In a joint histogram approach using equi-width bins, most bins are hardly populated. In other words, this representation of the data is highly inefficient. One way to address this issue is to perform an adaptive binning approach that partitions data recursively to generate *uniformly populated* bins rather than equi-width bins, as was done by Orlova et al. [4]. However, while this can potentially mitigate the issue of sparsity, the issue of binning granularity remains. Histograms that are too finely binned may be too sparse to reveal significant differences when compared. On the other hand, overly-coarse binning sacrifices resolving power. Hyper-rectangular histogram binning methods, whether equi-width or adaptive, tend to have limited success achieving a balance between data representation efficiency and resolving power [5].

An alternative representation of multidimensional distributions that optimizes efficiency and resolving power involves the use of “signatures” and “weights” rather than equi-width or adaptive histograms [5]. Signatures are defined as the main clusters (high-density regions) of a multidimensional distribution, and weights represent cluster size. This representation of images was found to yield the best results when comparing images to one another for the task of image retrieval. An analogy can be made to single-cell data modeling for the purpose of comparing single-cell samples, as single-cell samples are similarly multidimensional distributions. Using the signatureand-weights architecture, a “signature” can be thought of as a distinct cell subpopulation (or “cell subtype” e.g. memory B-cells or CD8^+^ effector T-cells), and the corresponding “weight” represents the number of cells in the cell subtype. The advantages of signatures and weights with respect to data representation efficiency may be intuitively extended from computer vision literature to our application; biologically relevant cell subtypes are the ideal signatures, or “bins,” for organizing single-cell data.

### Supplementary Note 3: Challenges of comparing multidimensional distributions represented as signatures and weights

Our final representation of a biological sample is a categorical frequency histogram representing the relative abundance (“weights”) of all possible cell subtypes (“signatures”) found in all samples collectively. Since our ultimate goal is to compare the similarity of samples, we need some metric to compare the similarity of these histograms. A major challenge is to identify a metric that captures the similarity of unique bins (i.e. “signatures” or “cell subtypes”) in the final distance measure. As a simple example using the EMT model, for a sample with 80% mesenchymal, 10% transitional, and 10% epithelial cells, we would expect a sample with 50% mesenchymal, 40% transitional, and 10% epithelial cells to be more similar (closer in distance) than a sample with 50% mesenchymal, 10% transitional, and 40% epithelial cells. This would be consistent with our intuitive sense of distance because 80-10-10 represents that most cells have fully transitioned from epithelial to mesenchymal states, 50-40-10 represents that most cells have partly or fully transitioned, and 50-10-40 represents that almost half of the cells have not transitioned at all. Earth Mover’s Distance is a distance metric that mathematically encodes this intuition and can be used as a robust measure of dissimilarity.

### Supplementary Note 4: “Tree-distance” as a measure of cross-bin dissimilarity

To more concretely explain our notion of “ground distance,” we will use our experiment including control and inhibited EMT samples as an example. What is the “ground distance” between the “bins” in this experiment? Recall that each bin represents a cell subtype (e.g. mesenchymal). Each cell subtype is associated with various different data points (individual cells assigned to that subtype), so it can be represented as the centroid of the cluster of cells that comprise it. Thus, we can quantify the ground distance between bins as the distance between their representative centroids.

To define a measure of distance between centroids, we first observe that, by design, all cells undergoing EMT across all samples originated from the same homogeneous epithelial population. Thus, it seems most reasonable to represent these cells as lying on a continuous trajectory with the epithelial cell subtype defined as the “origin” cell state. While EMT may have a primary linear progression from epithelial to mesenchymal state, we expect additional terminal cell states (e.g. apoptotic, senescent, proliferative) in our aggregate cell population. In other words, we expect the trajectory to be branched. In fact, many single-cell experiments represent data that are modeled well by branched trajectories, such as models of cell differentiation, cellular reprogramming, and immune response. To represent and visualize our data as a branched trajectory, we use Monocle 2, a tool for single-cell data that uses reversed graph embedding specifically for this purpose.

With the Monocle 2 embedding, we are able to visualize relationships between cell subtypes. In addition to providing a graph of all cells and a low-dimensional embedding that may be visualized, Monocle 2 also assigns each cell a “pseudotime” value to each cell: cells with a pseudotime of zero are in the starting epithelial state, while cells with high pseudotime values are further along the transition process (i.e. further from the starting epithelial state). This representation of our data as a graph embedding lends itself well to an intuitive sense of distance between any two cells: namely, the distance between the cells on the graph embedding’s minimum-spanning-tree (“tree distance”).

To compute tree distance between two distinct cluster centroids, we first map each centroid to the cell in its respective cluster that is geometrically closest (i.e. centroid of cluster A is represented by the cell in cluster A that is of the least Euclidean distance away). We then use the minimumspanning-tree of the graph constructed by Monocle and compute the shortest path between the two cells representative of their respective clusters. Finally, using pseudotime (PT) as a distance measure, we define the tree-distance between *cell*_1_ and *cell*_2_ as the following:

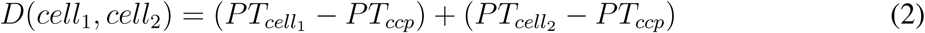

*ccp* represents the closest common progenitor of *cell*_1_ and *cell*_2_ and is defined as:

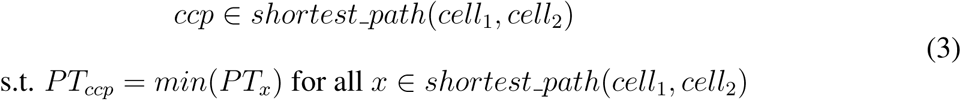

The shortest path between *cell*_1_ and *cell*_2_ is obtained from the minimum spanning tree modeled by Monocle 2 (Suppl. Fig. 2).

### Supplementary Note 5: Comparing EMD to existing methods of comparing two single-cell experimental conditions

Given the recent advent of multi-sample single-cell experiments, there are few methods designed to compare two experimental conditions measured at single-cell resolution. To date, there are two published methods suitable for this task: cellAlign and scUnifrac. In this section, we explore the properties and limitations of each in comparison to PhEMD’s approach to computing pairwise distances between experimental conditions.

cellAlign is a method that compares two experimental conditions (i.e., two heterogeneous cell populations) by first modeling each condition as an unbranched trajectory of cells, then assigning a pseudotime value to each cell based on its ordinal position in the trajectory, and finally computing a distance between the two experimental conditions as the “cost” of aligning the two pseudotemporal trajectories. A fundamental limitation of this approach is that it assumes all cells in an experimental condition can be accurately modeled as forming an unbranched trajectory. In reality, many experimental conditions consist of a heterogeneous mixture of cells that are best modeled as a branched trajectory or set of distinct clusters (i.e., distinct cell subpopulations). For example, in a simple set of cells consisting of CD4+/CD8+ double-positive (DP) T-cells differentiating into either CD4+ T-cells or CD8+ T-cells, the most correct model of all cells is a branched trajectory. Moreover, the pseudotime (PT) assignments most consistent with the underlying biology would be such that *PT* (DP T-cells) *< PT* (CD4+ T-cells) and *PT* (CD4+ T-cells) *≈ PT* (CD8+ T-cells). However, the assignment of the same pseudotime value to cells of different cell subtypes (as implicit when assigning pseudotime values to cells in branched trajectories) is not compatible with cellAlign, hence limiting cellAlign’s utility for analyzing such datasets.

sc-UniFrac is a different method of comparing two single-cell experimental conditions that works by aggregating all cells across all experimental conditions, performing hierarchical clustering of all cells, assessing the distribution of cells in each experimental condition across the cell clusters, and then computing a weighted UniFrac distance between conditions. A limitation of this approach stems from the fact that UniFrac distances are inherently expensive (resource-intensive) to compute. We found that sc-UniFrac faces scalability issues, as its memory requirements exceeded that of a standard laptop (2.5 GHz Intel Core i7 processor, 16 GB RAM) when attempting to compare experimental conditions containing collectively greater than 40,000 cells using default parameters. Another limitation of sc-UniFrac is that when used to perform pairwise comparisons of a set of more than two experimental conditions, sc-UniFrac returns only pairwise distances between conditions and offers no information on the source of differences (e.g., cell subpopulational differences) between experimental conditions. These limitations may be due to the fact that sc-UniFrac was primarily designed to be used to compare two experimental conditions rather than to perform pairwise comparisons of a set of many experimental conditions. Nevertheless, sc-UniFrac’s limited scalability and biological interpretability prevent it from being useful for analyzing large multi-sample datasets such as our drug-screen experiment spanning 300 experimental conditions and over 1.7 million cells.

Our implementation of EMD addresses the above limitations of both cellAlign and sc-UniFrac. By using Monocle 2 and the notion of “tree-distance” to relate cell subpopulations, we are able to model the cells as lying on a potentially branched manifold. Additionally, by explicitly modeling the global landscape of the cell-state space and representing each experimental condition as its percent abundance of each cell subtype, we are able to observe not only the degree of difference between conditions but also the specific cell subpopulations that are driving these differences. Finally, our approach is highly scalable and thus able to be applied to large datasets such as our newly generated drug-screen experiment. In contrast to sc-UniFrac, which was unable to be run on a laptop to analyze a set of 40,000 cells from two or more experimental conditions, PhEMD was successfully run on the same laptop to analyze a set of over 360,000 cells from 60 experimental conditions in under 15 minutes.

### Supplementary Note 6: Using Earth Mover’s Distance to construct an experimental variable state space embedding

We regard the analysis of data consisting of multiple samples as an extension of high-dimensional data analysis; in this case each sample is not only measured via many features, but also consists of many data points. The analysis of high-dimensional data, especially in unsupervised or exploratory settings, often introduces various challenges that are collectively referred to as the curse of dimensionality [6, 7]. A popular approach to analyzing such data is to use manifold learning methods that assume the intrinsic geometry of the data can conceptually be modeled as a low dimensional manifold (i.e., a collection of smoothly varying locally low dimensional data patches), which is immersed in the high dimensional ambient space of collected features [8]. Such methods often aim to uncover this intrinsic geometry by first capturing local neighborhoods, then using them to form a rigid structure of nonlinear relations in the data, and finally embedding this structure in low dimensions via a new set of features that preserve those relations (e.g., as distances).

At the core of most manifold learning methods is the assumption that there exists some natural distance metric that can be used to define local neighborhoods in the data. Indeed, popular manifold learning methods are often based on selection of nearest neighbors over simple Euclidean distance, even though this distance is only meaningful locally due to the curse of dimensionality. In multisample data, on the other hand, samples are no longer individual vectors, but rather they form data clouds with varying numbers of data points (i.e., cells). Therefore, to construct an intrinsic data geometry between samples, we have to first define (and compute) a notion of distance between samples, which can then be used for further analysis.

To compare two samples, we consider two notions of quantifying the difference between the distributions represented by them. First, two distributions can be compared by considering how distinguishable they are from each other. Indeed, if they are nearly indistinguishable from sampled data then the two samples should be considered very similar, while the easier it is to set their distributions apart, the more different the samples are from each other. This notion is typically considered in machine learning for generative tasks, e.g., to produce artificial images that are indistinguishable from real ones [9]. Second, two distributions can be compared by quantifying how hard it is to transform one distribution to another. If only a small perturbation is required then the samples are close together, while drastic changes mean they should be far apart. Remarkably, these two notions are closely related via the Kantorovich-Rubinstein duality theorem [10, 11], and can both be computed by the Earth Mover’s Distance (also known as Wasserstein metric) discussed in this work. Once the distance metric is formulated, the construction of an intrinsic manifold geometry from it amounts to computing pairwise distances between samples, organizing in a distance matrix, and passing this matrix as input to manifold learning methods, such as the diffusion maps [12] method used in this work, in order to construct a manifold geometry as described above.

### Supplementary Note 7: Leveraging all cells using Monocle 2 with nearest-node mapping

To ensure that all unique cell subtypes across all inhibition conditions were detected, we sampled cells from all inhibition and control conditions. Unique cell subtypes were then identified by running Monocle 2 on an equal subsampling of 200 cells from each condition. Through this step, each of the 200 subsampled cells from each inhibition condition was assigned to a specific subtype. Note that cells that were not initially subsampled still lacked a subtype assignment. Ideally, Monocle 2 would be performed using all cells from all inhibition conditions to assign subtypes to all cells. Unfortunately, this was not computationally tractable and raises the issue of scalability for Monocle 2 or alternative manifold-building algorithms that order and cluster high-dimensional data.

Our solution was to incorporate all cells into our analysis by iterating through the entire set of over 200,000 cells (including cells not initially subsampled) and mapping each to its most similar cell subtype. Note that each cell subtype detected by Monocle was defined as a cluster of cells with similar features. To assign cell *x*, which was not initially included in the construction of the Monocle cell-state embedding, to a cell subtype, we first identified cell *y* in the initial embedding that was most similar to cell *x*, i.e. the cell with the lowest Euclidean distance from cell *x*. Cell *x* was then given the same cell subtype assignment as cell *y*. The end result was that each cell was assigned to a specific cell subtype.

### Supplementary Note 8: Leveraging all cells using Seurat with nearest-node mapping

Similarly to our approach described in Supplementary Note 6, we ensured that all cell subtypes across all single-cell samples were represented in our cell state embedding by taking a random, equal subsample of cells from each sample. Louvain was then applied to the (batch-corrected) subsampled data, resulting in each subsampled cell being assigned to a cell subtype. To determine the cell subtypes of cells not initially subsampled, we used a nearest-neighbor approach similar to that described in Supplementary Note 6. For each cell *x* not initially assigned a cell subtype, we first identified the set of all cells *C* from the same batch that were assigned to a Louvain-based cell subtype. We then identified cell *y ∈ C* with the lowest Euclidean distance from cell *x* in the original protein expression space. Cell *x* was assigned the same cell subtype assignment as cell *y* for all cells x to ensure all cells were assigned a cell subtype.

### Supplementary Note 9: Characterizing EMT inhibitor groups identified by partitioning PhEMD inhibitor embedding from a single CyTOF run

Inhibitor Groups B-G represented inhibitors in which EMT was progressively less-strongly inhibited (Fig. 4). Group B represented 2 EGFR inhibitors that strongly halted EMT, as indicated by a marked predominance of epithelial cells at time of measurement. Group C represented an mTOR/PI3K inhibitor that had particularly low cell yield (Supplementary Table 5) and predominance of early transitional (C-2) and apoptotic (C-11) cells. Group D included 3 inhibitors that halted EMT moderately, with a predominance of epithelial (C-1), early transitional (C-2), and hybrid EMT (C-10) cells. Group E contained four HER2/EGFR inhibitors that generated a mixture of epithelial (C-1), transitional (C-2 through C-5), and mesenchymal (C-7) cells. Group F included two PI3K inhibitors that resulted in a predominance of transitional cells. Group G included five inhibitors that weakly inhibited EMT, consisting mostly of late transitional, mesenchymal, and hybrid EMT cells. Group A represented unperturbed controls and the TGFβ-receptor inhibitor condition, consisting almost entirely of epithelial cells. Group H represented uninhibited controls and inhibitors ineffective at halting EMT, consisting mostly of late transitional, mesenchymal, and hybrid EMT cells. Group I consisted mostly of Aurora kinase inhibitors and is discussed in greater detail in subsection ‘Effect of drug-inhibitor perturbations on the EMT landscape in breast cancer’ of the ‘Results’ section of the main text.

### Supplementary Note 10: Assessing batch effect in multi-run experiment

Batch effect is a well-known problem when comparing data from multiple single-cell RNA-sequencing [13,14] or CyTOF [15] experiments. Because of this, single-cell samples are ideally processed and measured in a single batch. However, comparing samples across experimental runs is still of great interest. In some cases, the sheer number of samples makes simultaneous processing impossible.

In other cases, the experimental design (e.g. time-series analysis) precludes sample processing on the same plate or gene profiling of all samples simultaneously. In order to enable these sorts of experiments, a number of methods have been recently published that correct for batch effect. We chose canonical correlation analysis (CCA), a new feature of the popular Seurat package, as our batch correction tool and demonstrated that PhEMD can leverage existing batch correction methods to compare hundreds of samples from 5 experimental runs.

To assess the presence of batch effect in our multi-plate experiment prior to batch effect normalization, we performed t-SNE dimensionality reduction on an equal, random subsample of cells from each batch (Suppl. Fig. 7). Since each batch used the same Py2T breast cancer cell line and contained a relatively similar mix of inhibition and control conditions, batches were expected to have more shared than non-shared cell subtypes. If true, this phenomenon would be appear as extensive inter-plate mixing in most regions of the t-SNE cell state space. This is because most sources of variation in the data were expected to be attributable not to the plate on which samples were cultured or CyTOF run in which samples were measured, but instead to sample-specific biology. Visualizing the t-SNE embedding and coloring cells by their original batch (Suppl. Fig. 8a), we noticed poor inter-plate mixing. This indicated that batch effect was present in the unnormalized data.

We then applied CCA to the expression measurements and ran t-SNE on the batch-corrected data (Suppl. Fig. 7b). Reassuringly, we noticed that there was strong inter-plate mixing when coloring cells in the t-SNE embedding by their original plate. This suggests that CCA effectively corrected for the technical sources of variation that appeared to be dominating the initial t-SNE embedding based on un-normalized expression data (Suppl. Fig. 7a). To assess whether batch effect correction not only removed technical sources of variation but also performed accurate data alignment, we examined the control conditions present on each plate. Two sets of identical control conditions were included on each plate: one set consisted of Py2T epithelial cells cultured with neither TGF-*β* nor drug inhibitor (”untreated controls”), and the other set consisted of Py2T cells stimulated with TGF-*β* and given no drug inhibitor (”uninhibited controls”). In our final clustering of samples, we found that all of the untreated controls from all 5 plates clustered together and consisted almost entirely of the same epithelial cell population. Similarly, all of the uninhibited controls from all 5 plates clustered together and consisted predominantly of late-transitional and mesenchymal cells. Moreover, inhibitors targeting the same molecular target tended to group together, irrespective of batch (e.g. Clusters D, E, F). These findings suggest that CCA accurately aligned the expression data.

## Supplementary figures

**Supplemental Figure 1:**
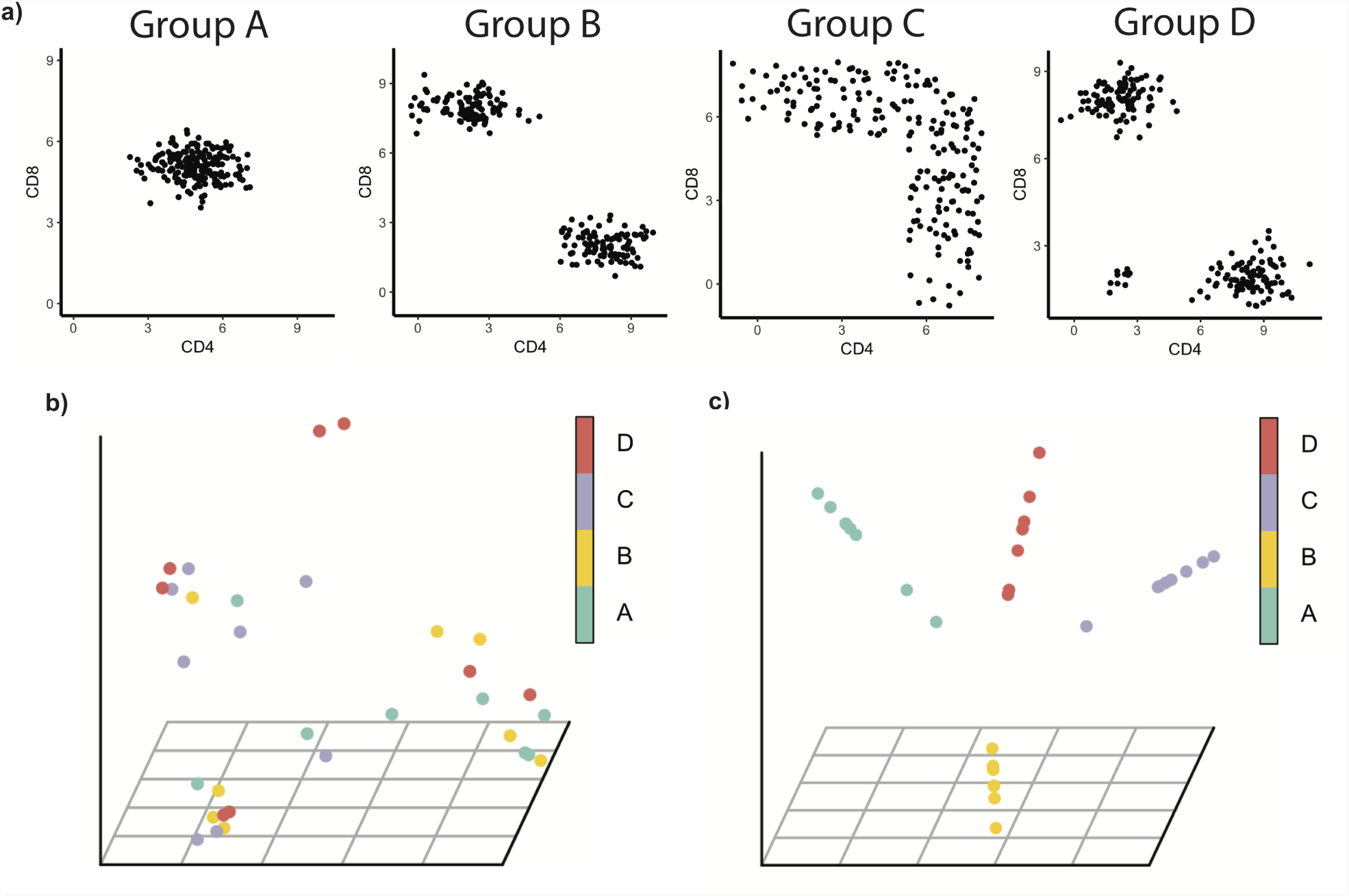
Single-cell analysis can resolve differences between biological samples that are indistinguishable on bulk (average) expression analysis. a) Single-cell profiles of each biological sample in a computationally-generated immune cell dataset. Groups A-D each had 8 samples that fit the single-cell profile. By design, all samples had roughly the same bulk expression of CD4 and CD8. b) Diffusion map embedding generated by embedding a sample-to-sample distance matrix, where pairwise distances between samples were computed by taking the Euclidean distance between samples represented as bulk expression of CD4 and CD8. Bulk expression profiles did not adequately reflect the biological differences between samples in this dataset and could not be used to distinguish samples in a biologically meaningful way. c) Diffusion map embedding generated by embedding a PhEMD distance matrix, which takes into account single-cell characteristics of each sample (see “Overview of PhEMD” in Results section). PhEMD successfully distinguished samples with different single-cell profiles from one another.

**Supplemental Figure 2:**
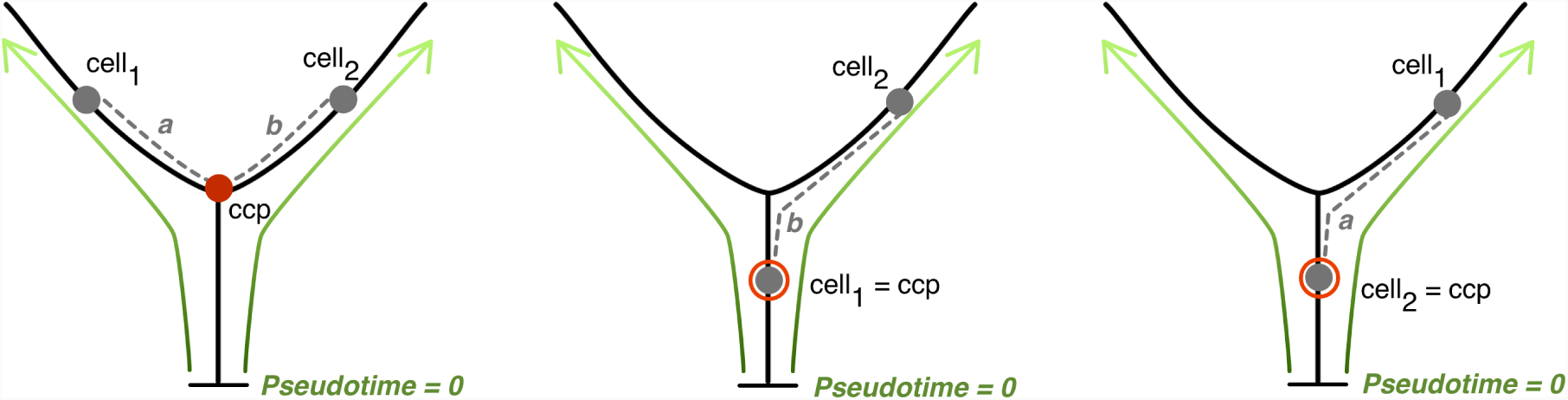
Visual representation of tree-distance computation for all possible relations between *cell*_1_ and *cell*_2_ on a Monocle 2 cell-state embedding. The dotted path *a* represents pseudotime distance from *cell*_1_ to *ccp* (closest common progenitor of *cell*_1_ and *cell*_2_) and *b* represents pseudotime distance from *cell*_2_ to *ccp*. Tree-distance between *cell*_1_ and *cell*_2_ is defined as *a* + *b* (see Supplementary Note 1).

**Supplemental Figure 3:**
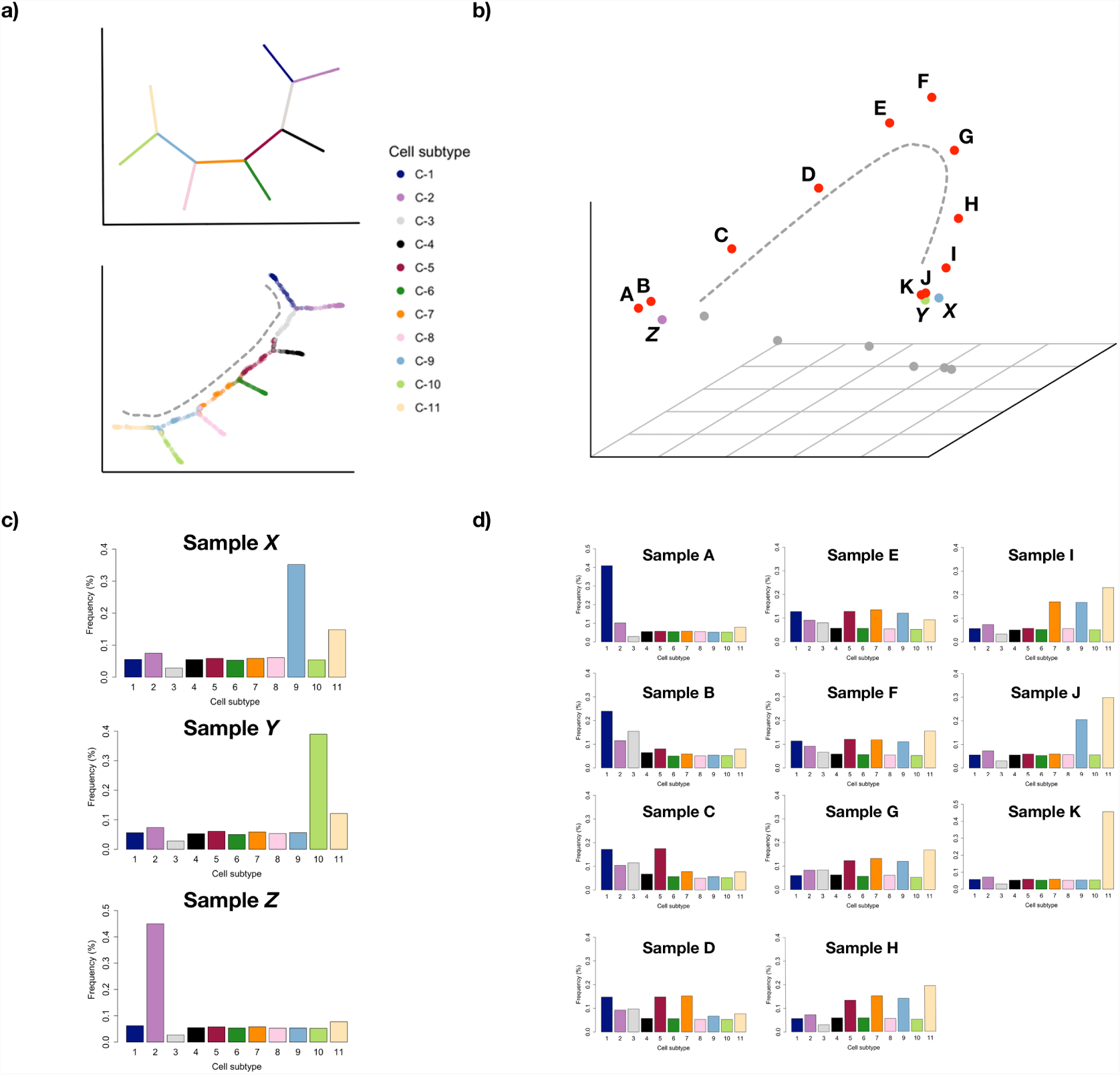
a) Ground-truth tree structure (top) and Monocle 2 embedding colored by ground-truth labels of cell-state space (bottom) for generated single-cell data. Grey dotted line denotes axis comprised of odd-numbered clusters (e.g. C-1, C-3,…) along which density is modulated for biological samples A–K. b) Diffusion map embedding of biological samples. Points colored red and labeled A–K represent samples that have density concentrated at various clusters along the trajectory from C-1 (“starting state”) and ending at C-11 (“terminal state”) highlighted in red. The alphabetical ordering of samples from A–K correspond to increasing intra-sample relative proportions of starting state to terminal state points. Samples X and Y represent samples with cells concentrated in clusters C-9 and C-10 respectively (i.e. highly similar cell subtypes), and Sample Z represents a sample with cells concentrated in cluster C-2 (highly dissimilar to cell subtypes C-9 and C-10). c) Relative frequency histograms representing distribution of cells across different “cell subtypes” (clusters) for Samples X, Y, and Z. d) Relative frequency histograms representing distribution of cells across different cell subtypes for Samples A–K.

**Supplemental Figure 4:**
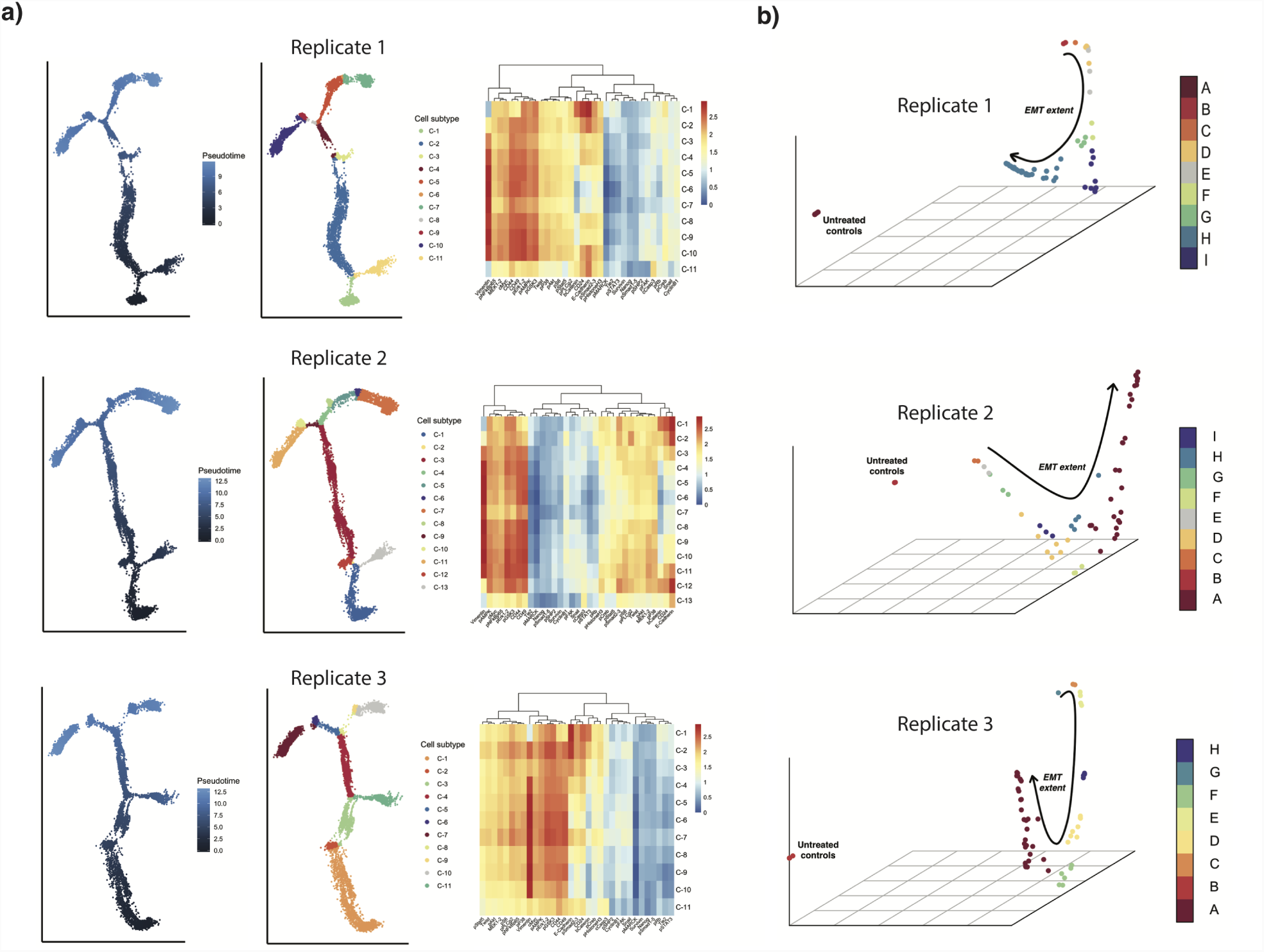
Figure demonstrating reproducibility of results across 3 biological replicates of our EMT data. a) Cell subtype expression patterns and cell state embeddings are similar across 3 replicates. b) PhEMD sample embeddings and inhibitor clusters are similar across 3 replicates (Supplementary Table 3).

**Supplemental Figure 5:**
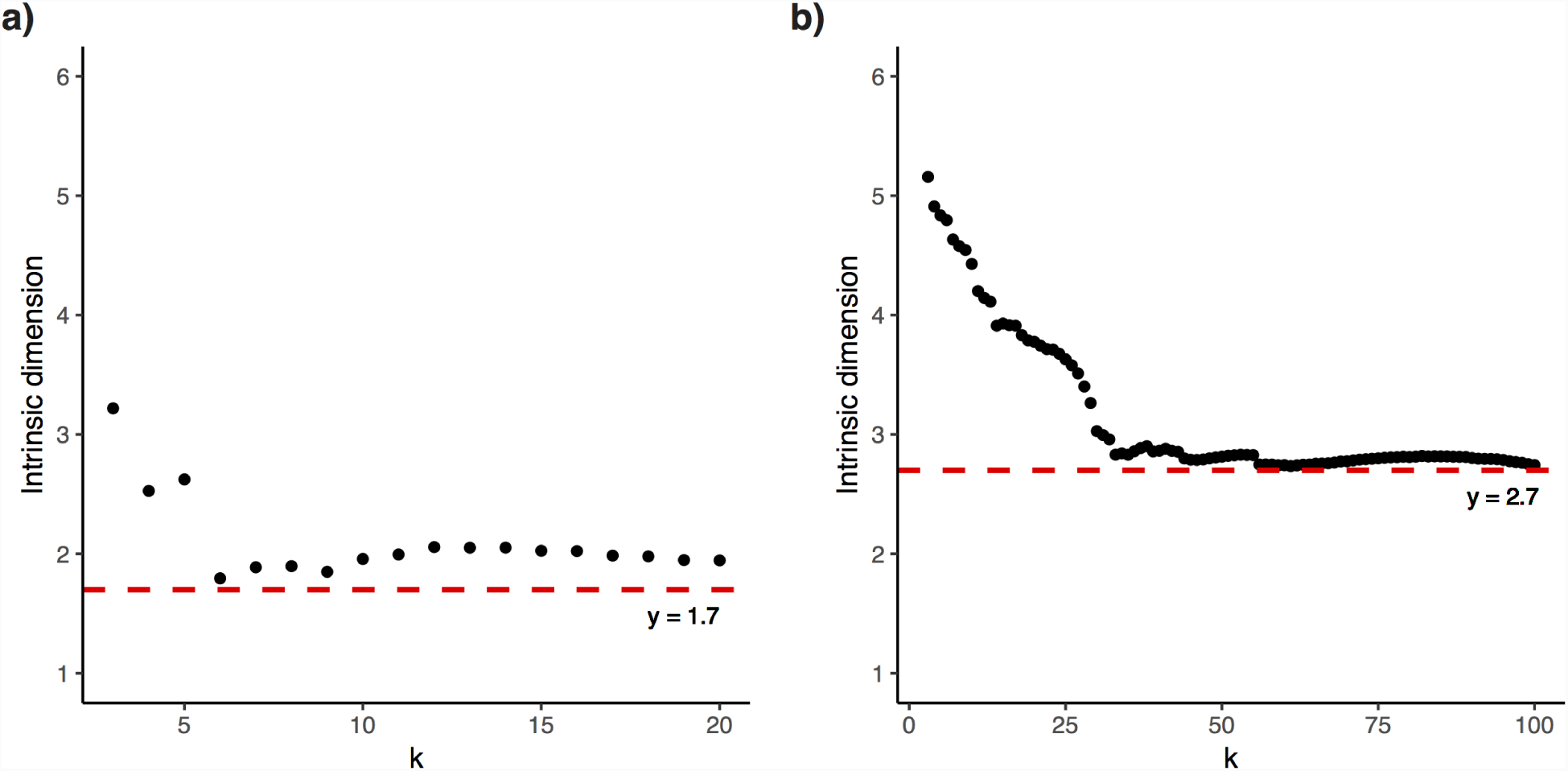
Intrinsic dimensionality analysis of the PhEMD embedding (i.e., network) of a) 60 single-batch and b) 300 multi-batch EMT inhibition and control conditions. Using the maximum likelihood estimation (MLE) approach, we found that intrinsic dimensionality was estimated to be 2 and 3 over a range of sufficiently large values of ‘k’ (knn parameter) for the single-batch and multi-batch embeddings respectively.

**Supplemental Figure 6:**
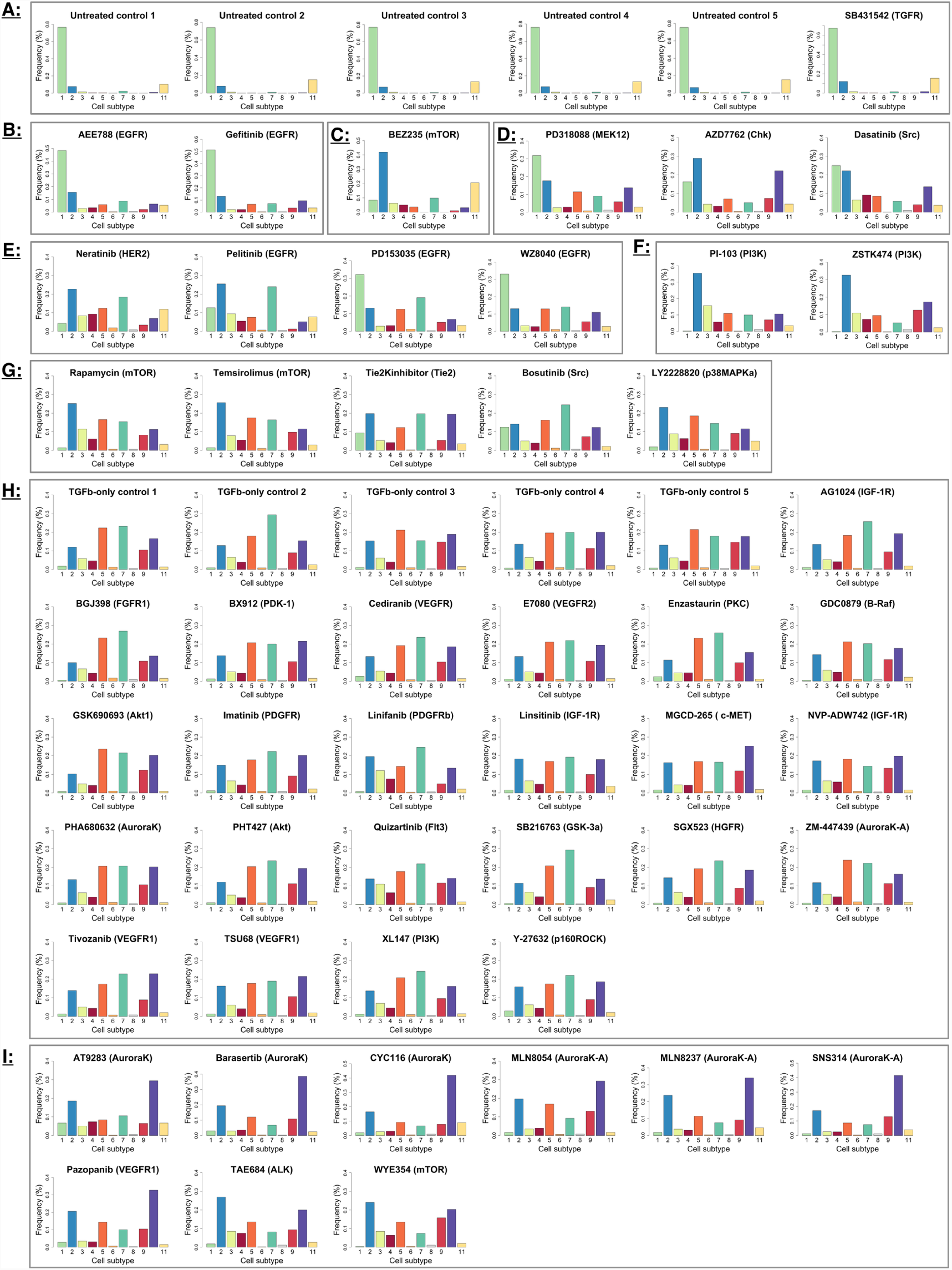
Frequency histograms representing distribution of cells across all cell subtypes in each inhibition condition of the EMT drug screen experiment. Cell subtypes are numbered and colored in accordance with the numbering and coloring of cell subtypes in Figure 4a. Letters denote final inhibitor groups, determined by hierarchical clustering of samples. Samples within each group demonstrate strong concordance with respect to cell subtype relative frequencies.

**Supplemental Figure 7:**
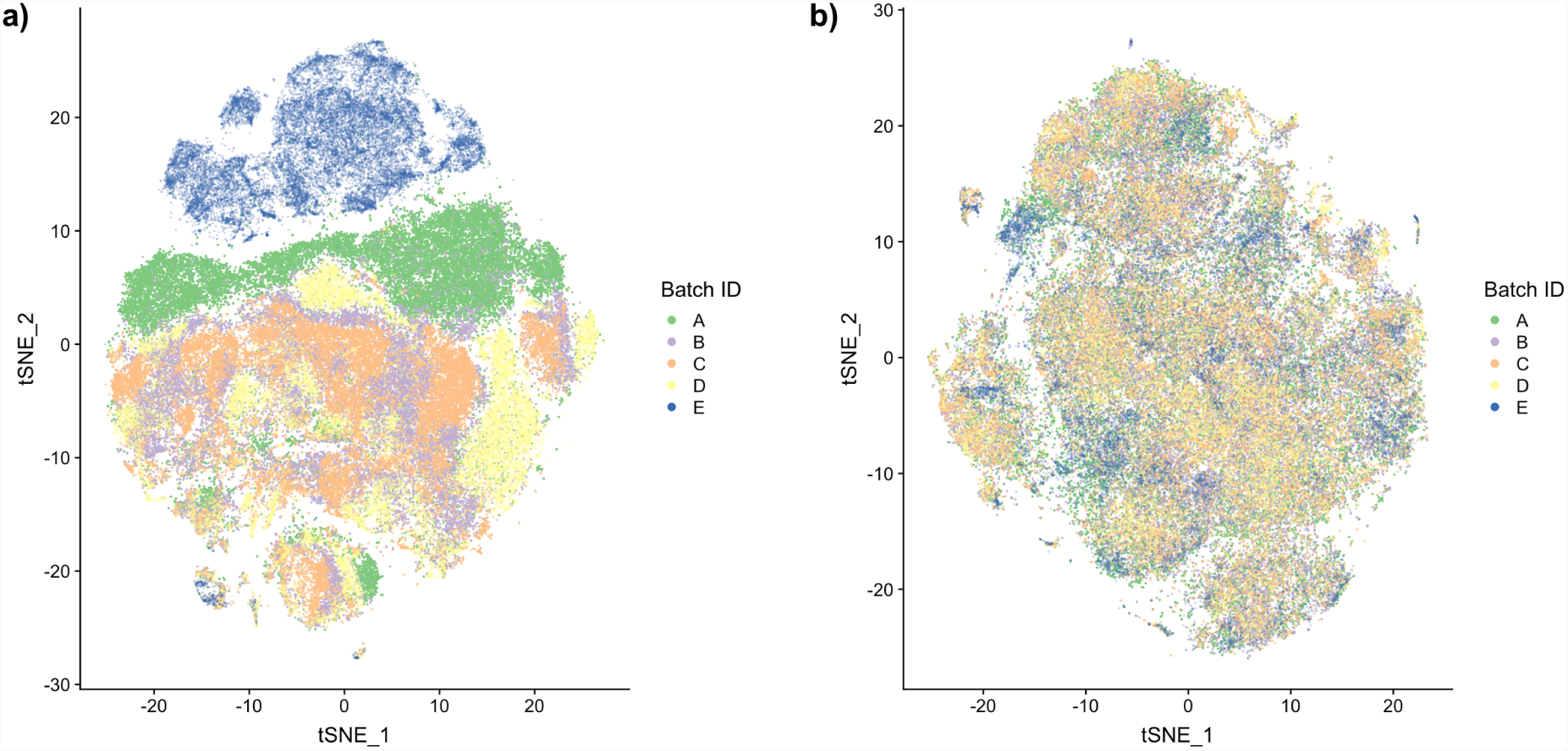
t-SNE clustering on cells from multiple CyTOF runs a) preand b) postCCA batch correction.

**Supplemental Figure 8:**
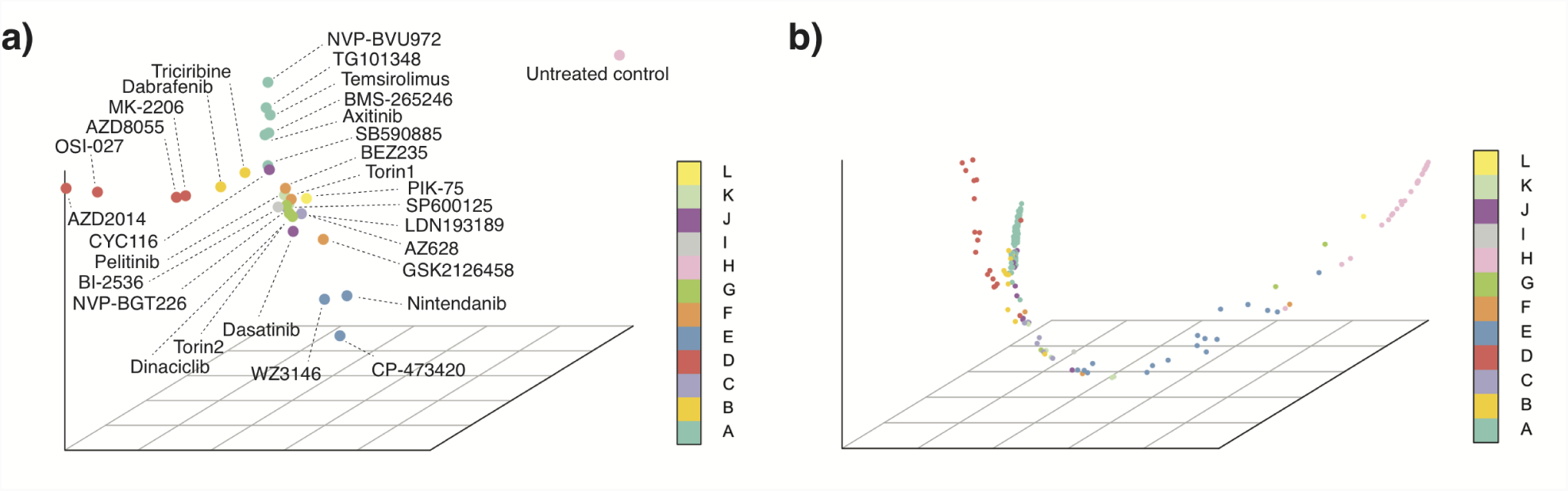
a) Diffusion map embedding of our 300-sample EMT experiment, plotting only the 30 landmark points identified using a previously published diffusion map sampling technique (see Online Methods). Points were colored based on cluster assignments as determined based on original clustering of all 300 samples (same as in Figure 5c). The 30 landmark points spanned all 12 inhibitor clusters identified in the original 300-sample embedding and visually captured the network geometry represented by the full 300-sample embedding (i.e., Figure 5c). b) Reconstructed diffusion map embedding by starting with the 30 landmark points and using a previously published out-of-sample extension technique to infer the embedding coordinates of all 300 samples relative to these 30 landmark (see Online Methods). The reconstructed diffusion map embedding closely resembled the original diffusion map embedding generated using pairwise distances between all 300 inhibitors (i.e., Figure 5c). This suggested that the 30 landmark points were sufficient to capture the full network geometry of the 300-sample experiment.

**Supplemental Figure 9:**
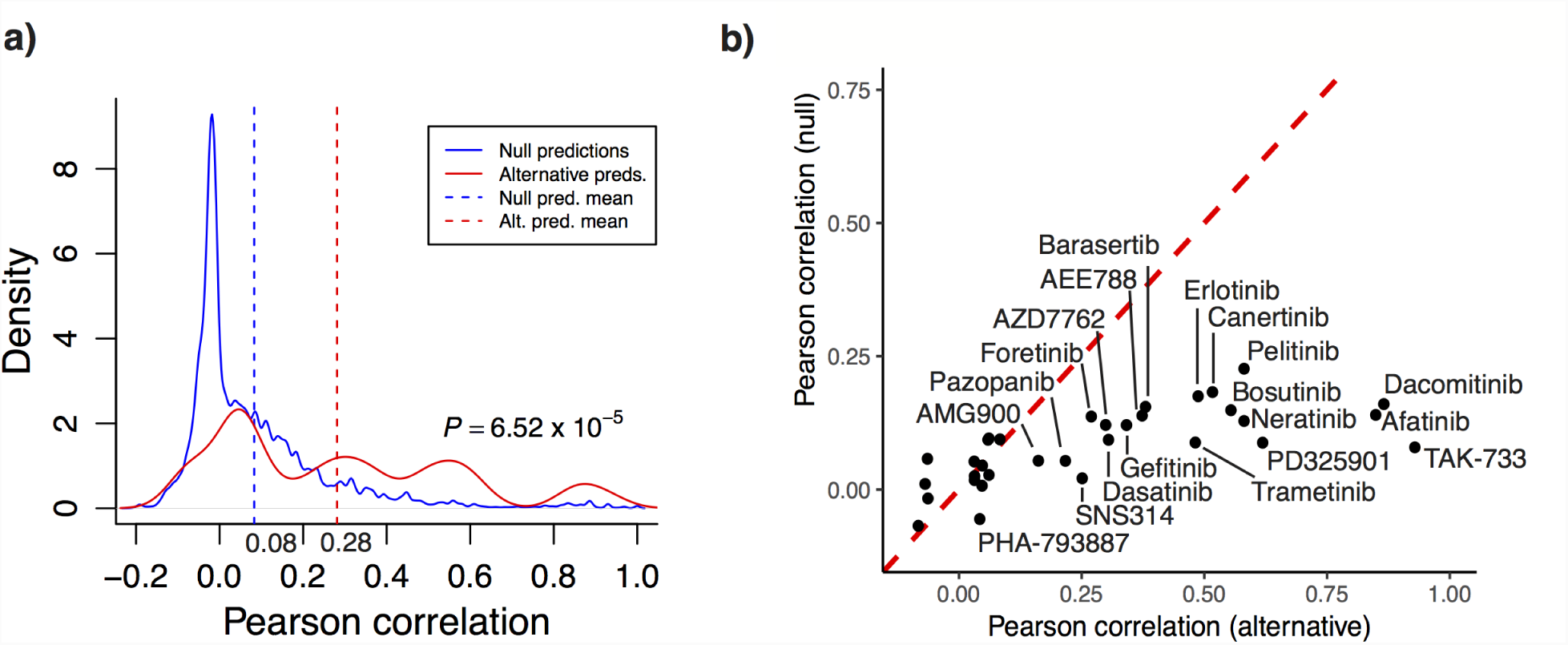
Assessing the utility of EMT perturbation screen PhEMD results for predicting known drug-target binding specificity. Prediction accuracy was defined as Pearson correlation between predicted and known drug-target binding specificity profiles. a) Probability density functions representing distribution of correlations between predicted and known drug-target binding specificity profiles. The null predictive model had poor prediction accuracy while the alternative model that incorporated PhEMD results from our EMT experiment performed significantly better (mean correlations of 0.08 vs. 0.28, *P*=6.52*10^−5^). b) Prediction accuracy of null vs. alternative models for predicting the drug-target binding specificity of each inhibitor. Given multiple null model predictions for each inhibitor, the y-axis represents the mean prediction accuracy of all predictions for a given inhibitor. See Online Methods for details on the null and alternative models presented.

**Supplemental Figure 10:**
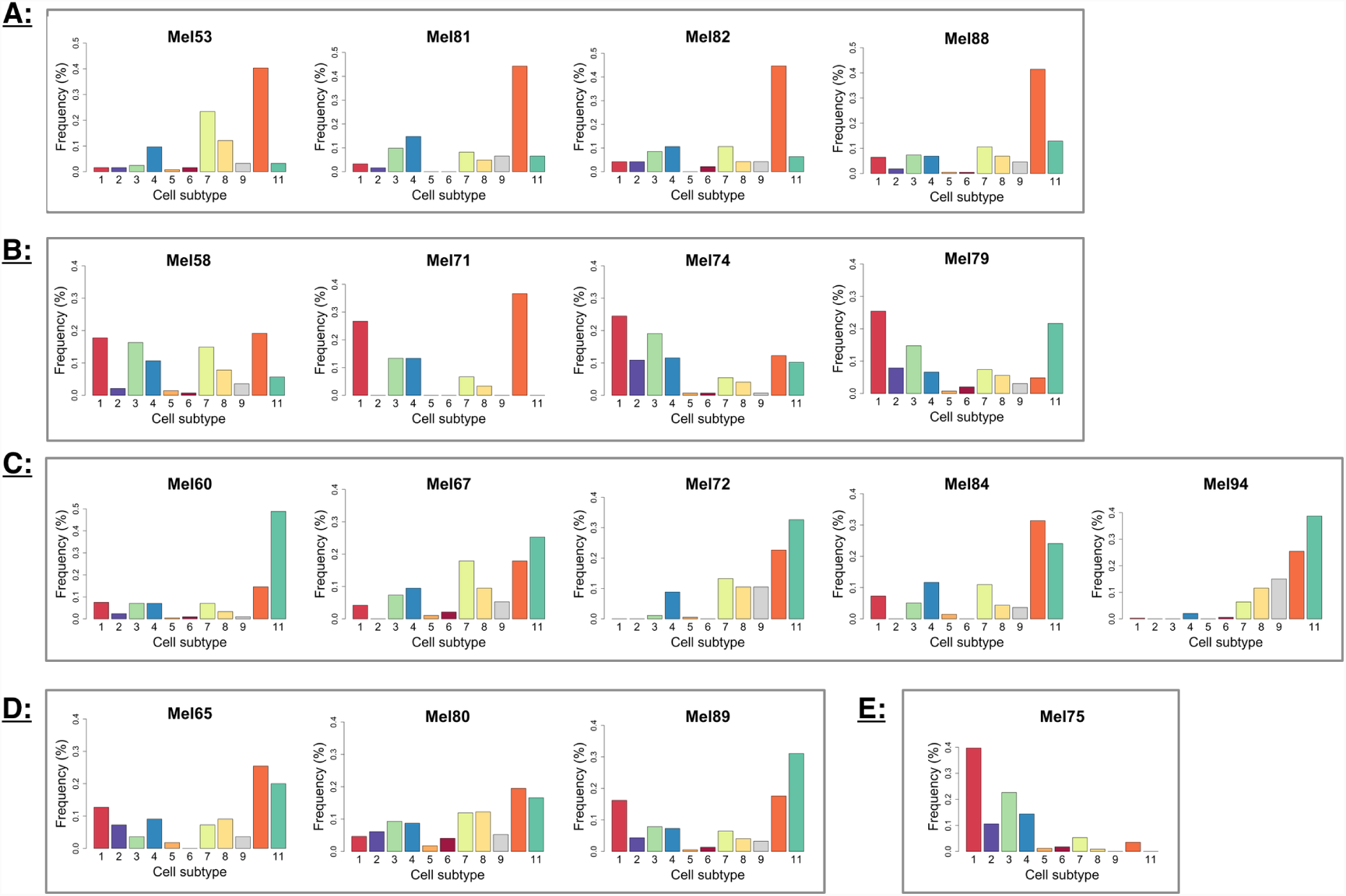
Relative frequency histograms representing distribution of cells across all cell subtypes in each tumor sample of the melanoma analysis. Cell subtypes are numbered and colored in accordance with the numbering and coloring of cell subtypes in Figure 5a. Letters denote final inhibitor groups, determined by hierarchical clustering of samples. Samples within each group demonstrate strong concordance with respect to cell subtype relative frequencies.

**Supplemental Figure 11:**
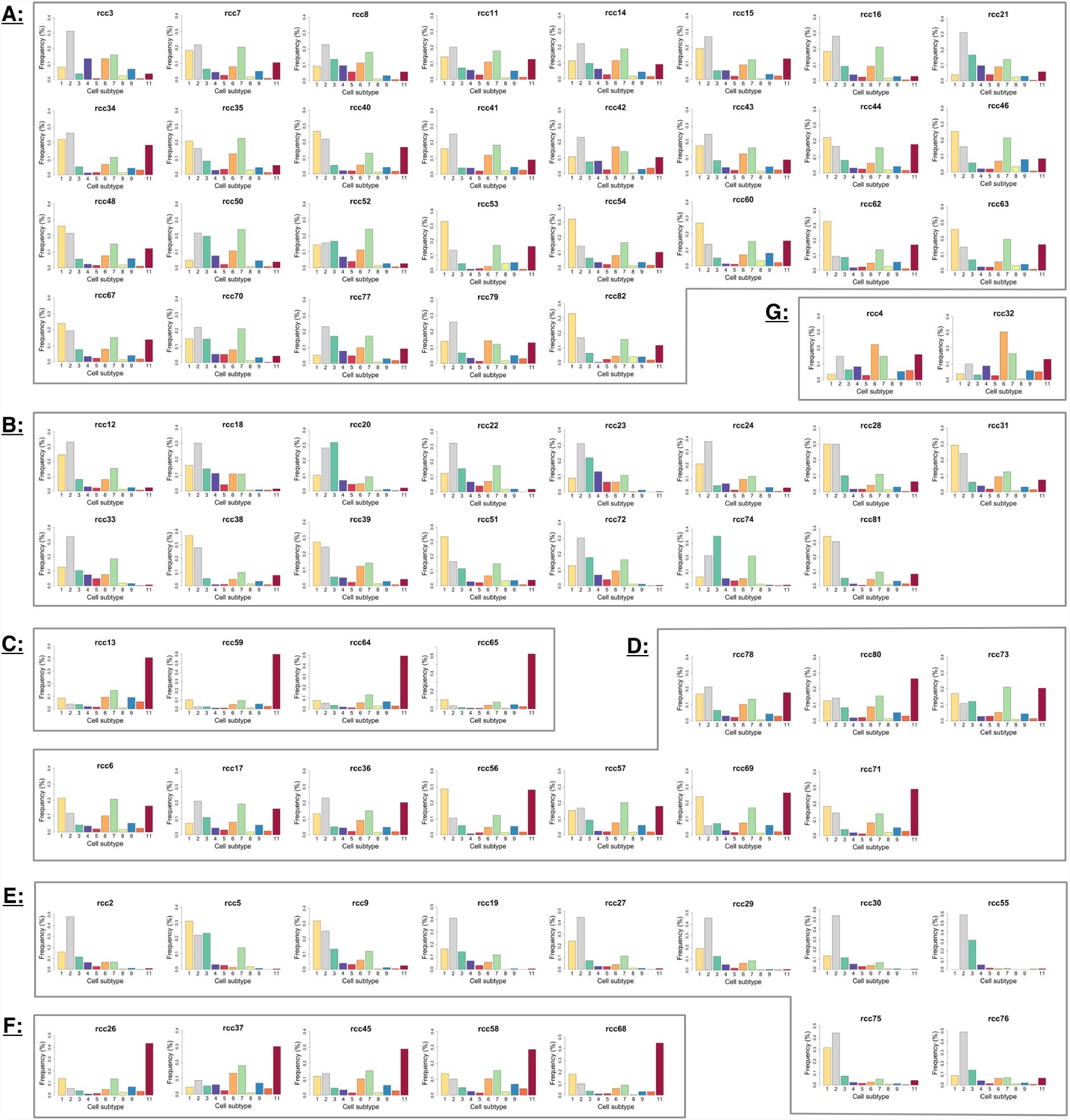
Relative frequency histograms representing distribution of cells across all cell subtypes in each tumor sample of the ccRCC analysis. Cell subtypes are numbered and colored in accordance with the numbering and coloring of cell subtypes in Figure 6a. Letters denote final inhibitor groups, determined by hierarchical clustering of samples. Samples within each group demonstrate strong concordance with respect to cell subtype relative frequencies.

